# Hepatocyte growth factor activator inhibitor-2 rapidly inactivates airway-expressed human Type II Transmembrane Serine Proteases

**DOI:** 10.64898/2026.01.23.701355

**Authors:** Bryan J. Fraser, Ryan P. Wilson, Jackie Lac, Olzhas Ilyassov, Almagul Seitova, Yanjun Li, Ashley Hutchinson, Yen-Yen Li, Peter Loppnau, Maria Kutera, Gregg B. Morin, Francois Benard, Cheryl H. Arrowsmith

## Abstract

Human Type II Transmembrane Serine Proteases (TTSPs) are essential entry factors for various influenza A and coronaviruses, and drive cancer metastasis when they are overexpressed by tumor cells. However, the natural inhibition mechanisms that regulate these proteases are not well understood. One natural transmembrane protease inhibitor, hepatocyte growth factor activator inhibitor-2 (HAI-2), has been shown to block TMPRSS2 activity and can prevent SARS-CoV-2 infection and reduce TMPRSS2-driven prostate cancer metastasis when overexpressed. In this study, we present biochemical and biophysical evidence showing that HAI-2 effectively inactivates TMPRSS2 and other TTSPs only after they have undergone zymogen activation. Through mutagenesis and ligand binding assays, we demonstrate that Kunitz Domain 1 (KD1) and KD2 can form stable ternary complexes with TMPRSS2 and other TTSPs, but do not employ the typical Laskowski inhibitor mechanism found for other macromolecular serine protease inhibitors. We also show that HAI-2 proteins do not inhibit the coagulation protease thrombin and that multivalent human IgG-tagged HAI-2 proteins are highly potent TMPRSS2 inhibitors. Our findings provide a mechanistic understanding of how TTSP activity is regulated in human airway cells and offer a foundation for developing engineered soluble HAI-2 proteins as anti-TTSP antivirals and anti-cancer therapeutics.

## Introduction

Human cell surface proteases play important roles in human biology and disease. A family of cell surface proteases, the Type II Transmembrane Serine Proteases (TTSPs), have been shown to have pivotal roles in epithelial homeostasis^1–3^, respiratory virus infections^4–13^, and cancer aggressiveness and metastasis^14–22^ through their serine protease activity. Accordingly, many TTSPs have presented as important drug targets for human diseases. TTSPs have a N-terminal single-pass transmembrane anchor and an ectodomain containing non-catalytic protein domains and a highly conserved S1 peptidase (SP) domain at their C-terminus that cleaves peptide and protein substrates containing Arg/Lys residues^23^. All TTSPs are initially expressed as inactive precursor proteins (zymogens) that require a proteolytic cleavage step to fully mature their SP domains and enable proteolysis (Fig. 1*A*). TTSPs can then go on to cleave other protease zymogens^16,24,25^, peptide hormones^26,27^, transmembrane growth factor receptors^17^, viral particles^4–6,9–13,28–32^, amongst other substrates before their activity is shut down by natural protease inhibitors^33–37^ and/or they are proteolytically removed from the cell surface (protease ectodomain shedding)^24,27,38^. Two TTSPs have emerged as important drivers of respiratory virus infections in humans, namely Transmembrane Protease, Serine-2 (TMPRSS2) and TMPRSS11D^4,5,9–11,13,30,39–42^. Cellular models of SARS-CoV-2 infection and Influenza A infection have shown that TMPRSS2 and TMPRSS11D rapidly cleave viral entry proteins, allowing virus-host membrane fusion that kickstarts viral infection. Our group has shown that TMPRSS2 and TMPRSS11D rapidly autoactivate, maturing their own proteolytic activity and could explain why they are preferentially exploited by coronaviruses for viral entry^43^. Furthermore, tool compounds that block TMPRSS2/11D activity prevent viral entry and pathogenesis in cellular and mouse models of SARS-CoV-2 and Influenza A infection^30,44,45^.

**Figure 1.**
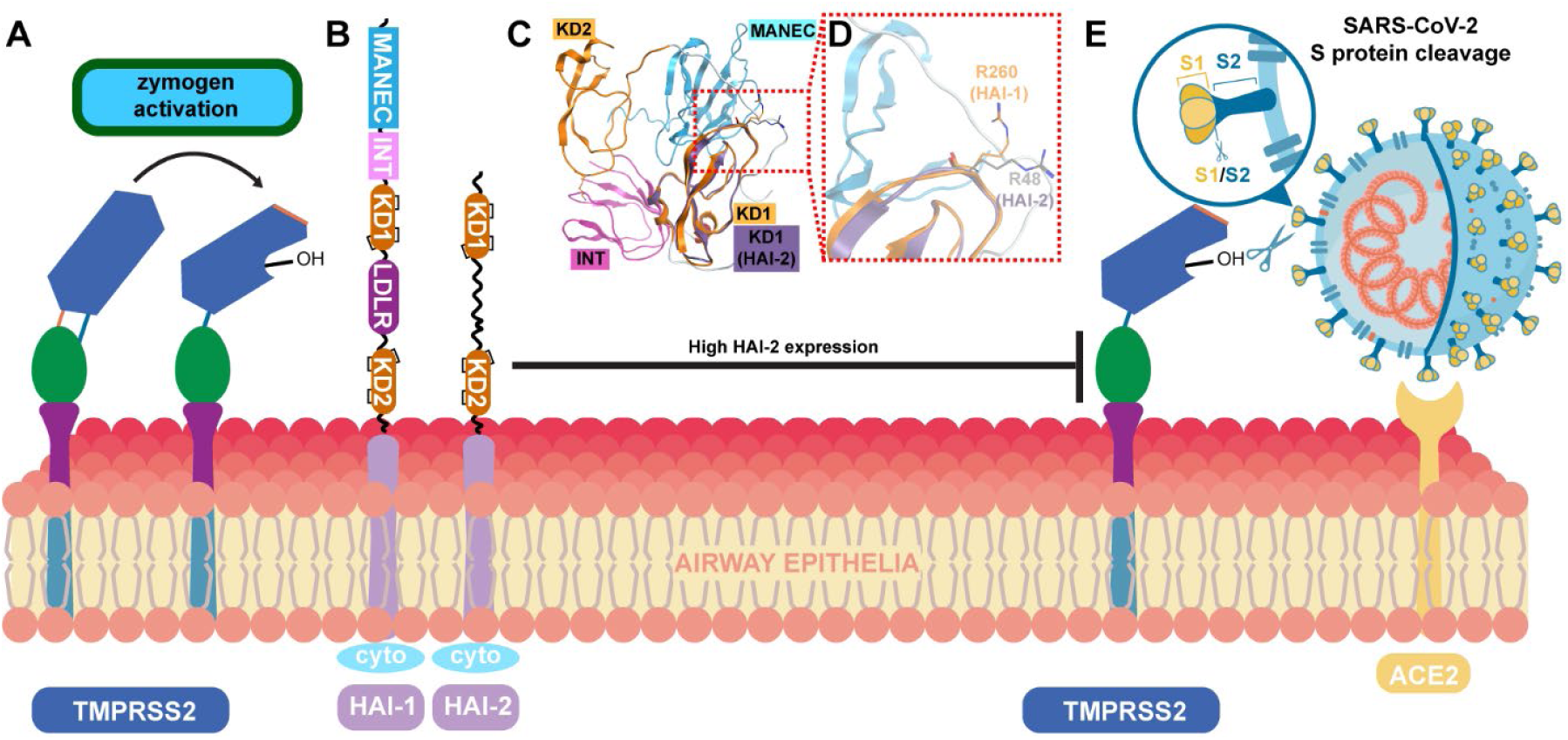
Cell surface TMPRSS2 undergoes zymogen cleavage activation and its protease activity is blocked by HAI-1 and HAI-2. **A**, Schematic of TMPRSS2 zymogen cleavage activation. Proteolytic cleavage at the TMPRSS2 R255-I256 peptide bond (shown as an orange stick) induces conformational changes that matures the TMPRSS2 catalytic domain and allows its catalytic S441 residue (shown as -OH) to cleave substrates. **B**, Schematics of the Hepatocyte growth factor Activator Inhibitor-1 (HAI-1) and HAI-2 proteins. MANEC-Motif At the N terminus with Eight Cysteines. INT-Internal PKD-like. KD1-Kunitz Domain 1. LDLR-Low Density Lipoprotein Receptor. KD2-Kunitz Domain 2. TM-transmembrane domain. Cyto – cytoplasmic domain. **C**, Cartoon representation of the crystal structures of the HAI-1 ectodomain and the HAI-2 KD1 domain. The KD1 domains of HAI-1 (orange cartoon) and HAI-2 (purple cartoon) are superposed and (**D**) show similar placement of their inhibitory Arg residues recognized by trypsin-like serine proteases. **E**, Schematic of TMPRSS2-driven SARS-CoV-2 viral entry of human airway cells. The SARS-CoV-2 viral particle docks to the human Angiotensin Converting Enzyme-2 (ACE2), bringing its Spike (S) protein in proximity to TMPRSS2 to enable S protein cleavage (shown as scissors).

Two key human proteins suspected to regulate TTSP activity are Hepatocyte Growth Factor Activator Inhibitor-1 (HAI-1; Uniprot ID O43278) and HAI-2 (Uniprot ID O43291)^20,46–49^. The HAIs are Type I transmembrane proteins that have been shown to inhibit the proteolytic activity of several TTSPs, including TMPRSS2, using their Kunitz Domains (KDs)^46,47,50,51^. HAI-1 and HAI-2 are broadly expressed across many tissue types but have slightly enhanced expression within the aerodigestive tract. Through KD binding to the catalytic domain of these proteases, HAIs prevent protease-mediated cleavage of substrates and the subsequent downstream signaling events. This inhibition is crucial for keeping the balance between proteolysis and anti-proteolysis in tissues, analogous to how serpins regulate the activity of blood serine proteases^52^. The reduced expression of HAI-2 or mutations that cause expression of dysfunctional HAI-2 protein have been linked to various disorders, including severe congenital diarrhea^53–55^, cancer progression^56–58^, and issues with spermatogenesis^59^. HAIs are located along the secretory pathway and at the plasma membrane, allowing them to block TTSP activity either during their trafficking to the plasma membrane or after they reach the plasma membrane^24,60^. However, previous studies have produced conflicting data on whether HAIs bind to TTSPs before or after zymogen activation to block TTSP protease activity^20,49,60,61^.

This work explores the molecular mechanism by which HAI-2 inhibits the protease activity of TMPRSS2 and other airway TTSPs implicated in respiratory virus infections and airway cancer aggressiveness and metastasis. We utilized engineered, soluble HAI-2 proteins to study the inhibitory and ligand binding potencies of the HAI-2 KD1 and KD2 domains against zymogen and active TTSPs. Through a combination of biochemical and biophysical assays, we demonstrate that HAI-2 specifically interacts with active TTSPs, not their zymogen forms. The HAI-2 proteins formed stable ternary complexes with airway TTSPs, inactivating their protease activities with comparable inactivation potencies to the small molecule ester nafamostat mesylate. Additionally, we showed that HAI-2 exhibits selectivity for airway TTSPs over other human trypsin-like serine proteases, such as thrombin. A chimeric HAI-2 protein produced in HEK293S cells with exceptional inhibitory potency towards TMPRSS2 is presented as a promising candidate for future anti-TTSP drug development campaigns for novel antivirals and cancer therapeutics.

## Results

### HAI-1 and HAI-2 proteins have a similar Kunitz Domain 1 but not Kunitz Domain 2

To understand molecular differences between the KD1 and KD2 domains of human HAI-1 and HAI-2, their sequences were aligned and the residues responsible for inhibiting trypsin-like serine proteases were compared (Fig. 2*A*). Most amino acid residues within the KD1 domains of HAI-1 and HAI-2 were identical or similar. Six cysteine residues participated in three nested, intradomain disulfide bonds (black bars; Fig. 2, *A* and *B*). The KD1 domain of HAI-1 and HAI-2 had respective protease inhibitory sequences of GRCR^260^GSFP and GRCR^48^ASMP (Position 1 (P1) arginine residues are numbered). Amino acid residues N-terminal to the P1 Arg are denoted as P2, P3, and P4, whereas residues C-terminal to the P1 Arg are denoted as Position 1’ (P1’), P2’, P3’, and P4’ (Fig. 1*B*). Thus, the protease inhibitory residues in the KD1 domains of HAI-1 and HAI-2 were only different at their P1’ and P3’ residues. The KD2 domain of HAI-1 and HAI-2 had respective protease inhibitory residues of GLCK^401^ESIP and GPCR^143^ASFP (Fig. 2, *C* and *D*). In contrast to the KD1 domains of HAI-1 and HAI-2, the KD2 domains of HAI-1 and HAI-2 showed more dramatic sequence differences within their protease inhibitory residues. The P4, P2, P2’, and P4’ amino acids within the KD2 inhibitory residue were identical, but the P3, P1, and P1’ residues were different. Notably, the KD2 domain of HAI-1 contained a P1 Lys residue and a P1’ Glu residue. The HAI-1 KD2 domain was shown to be incapable of blocking matriptase protease activity, whereas the HAI-1 KD1 domain could potently block matriptase protease activity^62^. Thus, the exact biological function of the KD2 domain of HAI-1 is unknown, and it is unclear if the KD2 domain of HAI-2 is similarly incapable of blocking TTSP protease activity.

**Figure 2.**
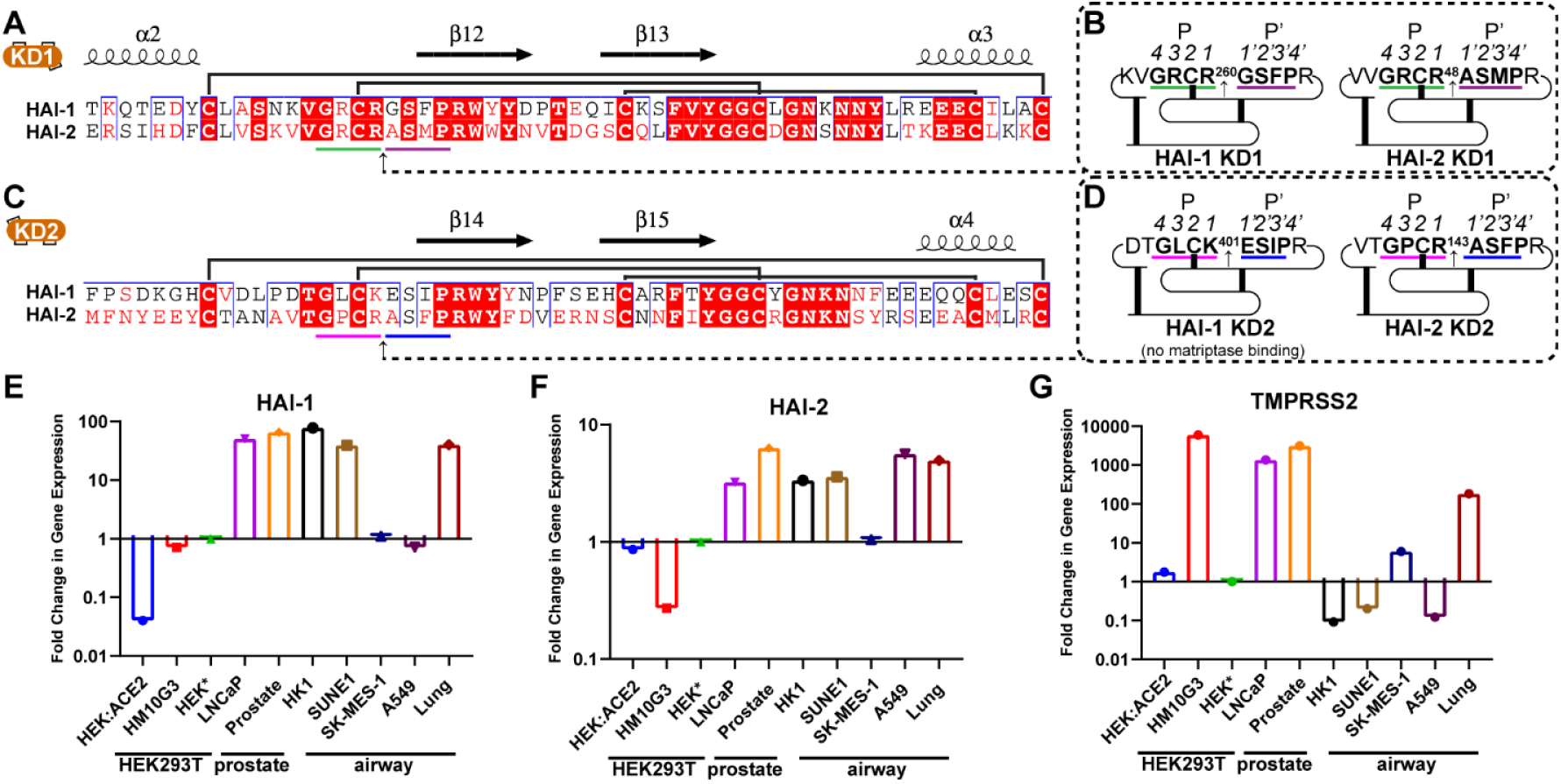
The HAI-1 and HAI-2 proteins have differences in their protease inhibition sequences that engage protease active sites. **A**, Sequence alignment of the KD1 domains of HAI-1 and HAI-2. An arrow denotes the scissile peptide bond of the KD1 domain where a trypsin-like serine protease would cleave a typical substrate. A green bar indicates the KD1 Position 1 (P1) arginine residue and neighboring P2-P3-P4 residues that also bind to trypsin-like serine binding sites S1-S2-S3-S4. A purple bar indicates the P1’-P2’-P3’-P4’ residues. **B**, Schematic of the 2D structure of the KD1 domains. Bolded bars indicate disulfide bonds that constrain the tertiary structure of the domain. **C**, Sequence alignment of the KD2 domains of HAI-1 and HAI-2 and (**D**) schematic of their 2D structures. **E**, HAI-1 cDNA levels quantified from the indicated cell lines or whole tissue prostate or lung RNA. Reverse transcriptase qPCR was used to measure relative mRNA transcript levels. Relative quantification of expression of the sample cell lines was preformed using the Livak method. A HEK293 cell line was set as a standard for relative HAI-1 quantification, indicated with an asterisk, *. Cell line details are available in Materials and Methods. **F**-**G**, HAI-2 and TMPRSS2 cDNA levels within the indicated cell lines.

### *HAI-1* and *HAI-2* are co-expressed with *TMPRSS2*

To verify that the HAI-1 and HAI-2 proteins are co-expressed with TMPRSS2, we quantified mRNA transcript levels of *HAI-1*, *HAI-2*, and *TMPRSS2* across various cell lines. HEK293T, prostate, and airway cell lines, as well as total human lung and prostate mRNA were tested, and expression levels were normalized to a standard HEK293T cell line (indicated as HEK*; Fig. 2, *E*-*G*). For HAI-1, transcript levels were low in HEK293T cell lines and in the epithelial non-small cell lung cancer cell lines SK-MES-1 and A549, but transcript levels were high in prostate and the upper airway cancer cell lines SUNE1 and HK1 (Fig. 2*E*). HAI-2 expression trends were similar to HAI-1, except for the A549 cell line where HAI-2 levels were high and HAI-1 levels were low (Fig. 2*F*). TMPRSS2 transcript levels were high in TMPRSS2-HEK293T cells, prostate cell lines, and in total human lung RNA (Fig. 2*G*). These data confirmed that HAI-1, HAI-2, and TMPRSS2 are expressed in the lung, and slight differences are apparent in the expression of these proteins for different airway cancer cell lines.

### Soluble HAI-2 KD1 domain protein rapidly inactivates TMPRSS2 activity

We designed and expressed soluble HAI-2 KD1 domain protein constructs for large-scale protein production and purification using the baculovirus expression vector system and Sf9 insect cells (Fig. 3*A*; *Fig. S1A*). The secreted KD1 domain protein was purified from media to homogeneity and produced a protein band at 12 kDa (Fig. 2*A*). KD1 potently inhibited TMPRSS2 activity with an IC_50_ of (7±3) nM, comparable to nafamostat’s potent TMPRSS2 inhibition with an IC_50_ of (2.1±0.2) nM (Fig. 2*B*). In contrast, 10 µM KD1 R48A mutant protein failed to inhibit TMPRSS2 activity(Fig. 2*B*), confirming R48 was critical for binding within TMPRSS2’s substrate binding cleft. To probe KD1’s mechanism-of-action, we used TMPRSS2 coaddition assays where substrate and inhibitor are simultaneously transferred to the enzyme. In this assay format, covalent inhibitors gradually disable TMPRSS2 throughout the assay, producing a plateau in enzyme activity (Fig. 2*C*). KD1 also inactivated TMPRSS2 activity over time (Fig. 2*D*). Through progress curve fitting, the TMPRSS2 inactivation potencies (*k*_inact_/K_I_) of KD1 was determined to be 0.21 s^-1^µM^-1^, comparable to nafamostat’s TMPRSS2 *k*_inact_/K_I_ value of 2.8 s^-1^µM^-1^ determined previously^63^.

**Figure 3.**
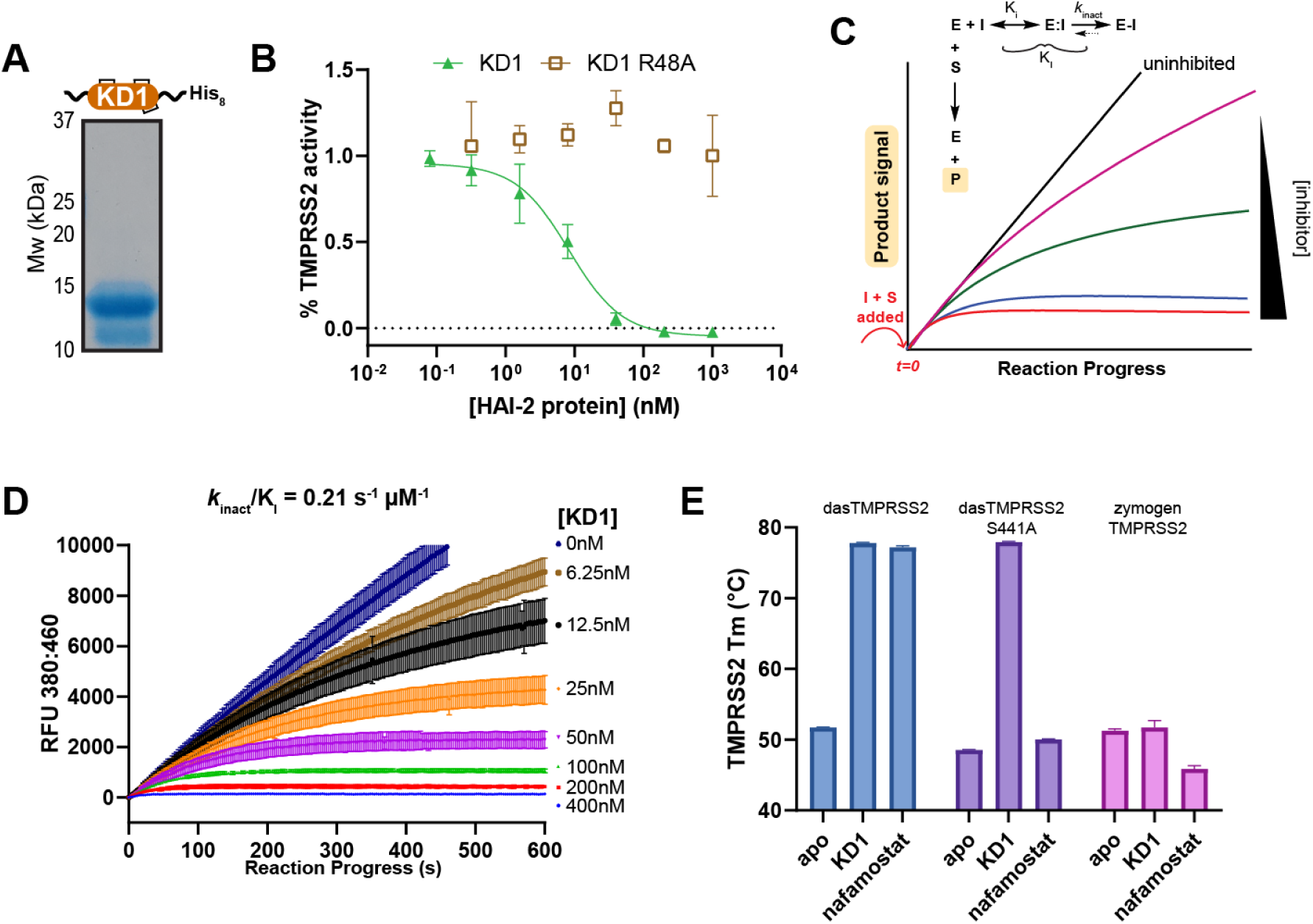
Soluble HAI-2 KD1 domain potently inactivates TMPRSS2 activity following zymogen activation. **A**, Purified, soluble HAI-2 KD1 domain. The His-tagged, secreted HAI-2 KD1 domain was purified through Ni^2+^-NTA IMAC followed by size-exclusion chromatography. **B**, TMPRSS2 half-maximal inhibitory concentration (IC_50_) plots with the KD1 and mutant KD1 R48A proteins. **C**, Schematic of a TMPRSS2 coaddition assay when substrate (S) and an irreversible protease inhibitor (I) are added simultaneously to the protease at *t=0* (red text). At increasing concentrations of irreversible inhibitor, the TMPRSS2 protease activity plateaus at a faster rate. **D**, Reaction progress curves for TMPRSS2 with the indicated concentrations of HAI-2 KD1 protein added to TMPRSS2 simultaneously with 100 µM Boc-QAR-AMC substrate. Progress curve fitting was performed using the DynaFit software to determine TMPRSS2 *k*_inact_/K_I_ values (Experimental Procedures). **E**, TMPRSS2 melting temperatures (T_m_s) in the absence (apo) or presence of the indicated ligands.

### KD1 inactivates TMPRSS2 activity through a nonstandard inhibition mechanism

Macromolecular inhibitors of serine proteases typically employ the Laskwosksi, or “Standard” inhibition mechanism where the protease uses its catalytic serine residue to nucleophilicially attack an appropriately positioned, scissile peptide bond (between P1 and P1’)^64–67^. The large covalent adduct excludes active site water molecules from completing the hydrolysis reaction, trapping the protease mid-catalysis. The protease:inhibitor complex then cycles through breakage and ligation of the scissile peptide bond, effectively preventing other substrate’s access to the protease. This mechanism-based inhibition strategy has been demonstrated for bovine pancreatic trypsin inhibitor (BPTI; apronitin)^68,69^, sunflower trypsin inhibitor-1 (SFTI-1), and Kazal-type inhibitors including serine protease inhibitor, kazal-type 1 (SPINK1)^70,71^. To investigate if the HAI-2 KD1 domain employs the Laskowski mechanism to inhibit TMPRSS2, we performed ligand bind assays with three forms of TMPRSS2; i) activated dasTMPRSS2^72^, ii) activated dasTMPRSS2 S441A, and iii) zymogen TMPRSS2 (Methods). By using biophysical binding assays rather than enzyme activity assays, we could measure the TMPRSS2:KD1 interaction in the presence or absence of TMPRSS2 catalytic activity. We opted to use differential scanning fluorimetry (DSF) to measure ligand-induced shifts in the TMPRSS2 melting temperature, a strategy previously applied for TMPRSS11D^73^. All three TMPRSS2 proteins produced typical melt curves across a temperature gradient of 20-95°C (detected by SYPRO orange fluorescence; Fig. S1A) and their Melting Temperatures (T_m_s) were determined to be (52.8±0.1)°C, (49.5±0.4)°C, and (51.3±0.2)°C for dasTMPRSS2, dasTMPRSS2 S441A, and zymogen TMPRSS2, respectively. In contrast, the KD1 protein did not produce a detectable melt curve in the DSF assay (Fig. S1B). For dasTMPRSS2 and dasTMPRSS2 S441A, the KD1 protein induced large TMPRSS2 Tm shifts (ΔT_m_s) of (24.8±0.1)°C and (29.4±0.1)°C, respectively, but no zymogen TMPRSS2 ΔT_m_ was detected (Fig. 2*E*). These data indicated that KD1 exclusively forms a complex with active TMPRSS2 (but not zymogen TMPRSS2), and the thermal stability of this complex does not depend on TMPRSS2’s catalytic serine residue. In contrast, nafamostat only induced a ΔT_m_ for active dasTMPRSS2 (ΔT_m_ = (24.4±0.2)°C) and not dasTMPRSS2 S441A, indicating that nafamostat primarily relies on the TMPRSS2 catalytic serine residue for complex formation and TMPRSS2 thermal stabilization, similar to TMPRSS11D^73^. Thus, the HAI-2 KD1 domain is a noncovalent inactivator of TMPRSS2 activity that does not rely upon classically understood, mechanism-based inhibition to achieve its nanomolar binding potency.

### Mutation of A49 modulates HAI-2 KD1’s mechanism of TMPRSS2 inhibition

To further probe how KD1 inactivates TMPRSS2, we mutated the P1’ residue of KD1, Ala49, to monitor how changes around the P1-P1’ peptide bond impact the inhibition mechanism. A total of 20 KD1 proteins (1 wild-type and 19 mutants) were produced and purified. Notably, the HAI-2 KD1 A49G mutant’s sequence (GRCR^48^GSMP) was nearly identical to the HAI-1 KD1 domain sequence (GRCR^260^GSFP). All KD1 mutant proteins were screened for TMPRSS2 inhibition at a KD1 protein concentration of 1 µM (Fig. S2B). Most HAI-2 KD1 mutant proteins showed no inhibition of TMPRSS2 peptidase activity, except wild-type KD1, A49G, A49S, A49T, and A49M which had TMPRSS2 IC_50_s of (11±2), (9±1), (14±4), (40±10), and (14±1) nM, respectively (Fig.4A). The wild-type KD1, KD1 A49G, KD1 A49S, and KD1 A49M proteins had TMPRSS2 *k*_inact_/K_I_ values of (0.21±0.02), (0.22±0.03), (0.15±0.03), and (0.025±0.003) s^-1^µM^-1^, respectively (Fig. S2C). TMPRSS2 reaction progress curves in the presence of the KD1 A49M protein showed an initial TMPRSS2 inactivation phase but did not completely plateau. This progress curve behaviour suggested that the KD1 A49M protein gradually dissociated from the TMPRSS2 substrate binding cleft and restored substrate access to the protease active site. TMPRSS2 reaction progress curves with KD1 A49T resembled progress curves typically produced in the presence of a competitive inhibitor like 6-amidino-2-naphthol (Fig. 4B, and C). KD1 A49T data was curve-fitted to determine a TMPRSS2 inhibition constant (K_i_) of 196 nM. Thus, modifications at the P1’ residue had dramatic implications for HAI-2 KD1 inhibitory potency and longevity of the TMPRSS2:KD1 ternary complex.

**Figure 4.**
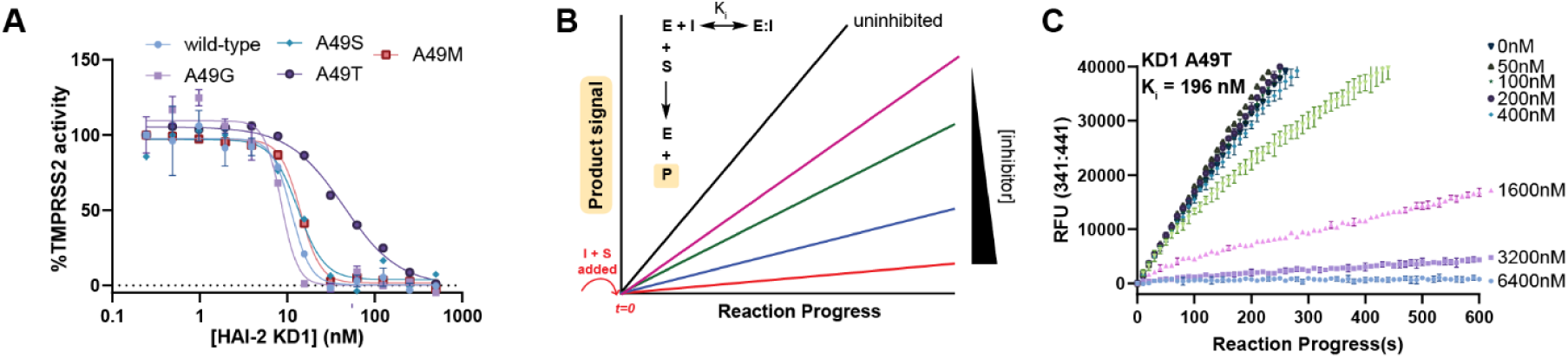
Mutations targeting the HAI-2 KD1 A49 residue impact TMPRSS2 binding affinity and mechanism-of-action. **A**, TMPRSS2 IC_50_ plots for the indicated HAI-2 inhibitor proteins in peptidase inhibition assays with 100 µM Boc-QAR-AMC substrate. Reaction velocities were tabulated across the first 60 seconds following substrate addition and relative TMPRSS2 activity was determined through normalization with an uninhibited TMPRSS2 sample. Datapoints are expressed as mean values +/- sd for *n=2* biological replicates with *n=3* technical replicates (*n=6* total) with consistent results obtained across *n=4* independent biological experiments. **B**, Schematic of a TMPRSS2 coaddition assay when substrate (S) and a reversible protease inhibitor (I) are added simultaneously to the protease at *t=0* (red text). **C**, Reaction progress curves for TMPRSS2 with the indicated concentrations of HAI-2 KD1 A49T protein added to TMPRSS2 simultaneously with 100 µM Boc-QAR-AMC substrate. Progress curve fitting was performed using the DynaFit software to determine TMPRSS2 K_i_ values (Experimental Procedures).

### TMPRSS2:KD1 complexes are stable to gel filtration and mild SDS treatment

To determine if the TMPRSS2:HAI-2 complexes would dissociate after a large dilution step, we evaluated if the TMPRSS2:KD1 protein complex could be isolated by size-exclusion chromatography on a Superdex 75 (S75) preparative grade 124 mL gel filtration column. Active TMPRSS2 was incubated with excess KD1 protein (protease:inhibitor ratio of 1:5) for 10 minutes prior to injection to the gel filtration column. Two absorbance peaks eluting at 59 mL and 89 mL corresponded to the TMPRSS2:KD1 complex and isolated KD1 protein, respectively (Fig. 5A). The 59 mL elution peak was loaded to a SDS-PAGE gel (sample prepared with SDS lacking 2-mercaptoethanol) and a prominent protein band migrated at ∼50 kDa (Fig. 5B), confirming the formation of a dilution-stable and SDS-stable TMPRSS2:KD1 complex. When the SEC chromatograms of the isolated TMPRSS2 protein and the TMPRSS2:KD1 co-injected sample were overlaid, a large shift in elution volume was observed between the TMPRSS2 sample (71 mL peak) and the TMPRSS2:KD1 sample (59 mL peak; Fig. 5*C*). The individual TMPRSS2 and KD1 proteins could be resolved by heating the sample prior to SDS-PAGE gel separation (Fig. 5D).

**Figure 5.**
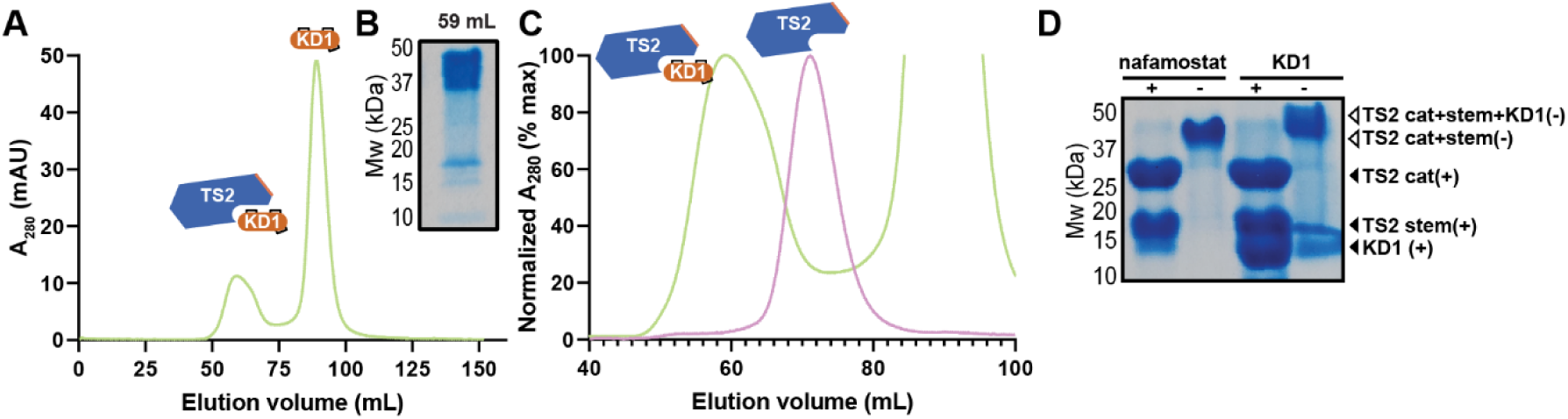
TMPRSS2:KD1 ternary complexes are isolatable by size-exclusion chromatography. **A**, Size-exclusion chromatography (SEC) chromatogram of the dasTMPRSS2:KD1 complex injected to a HiLoad 16/60 Superdex 75 124 mL gel filtration column. The two absorbance peaks are labelled as the TMPRSS2:KD1 complex and isolated KD1 protein. **B**, OThe 59 mL elution fraction was separated on an SDS-PAGE gel. The SDS-PAGE sample was prepared without reducing agent and was not thermally denatured in advance of gel separation. **C**, Overlaid SEC chromatograms of nafamostat-treated TMPRSS2 (pink trace) and TMPRSS2:KD1 (green trace). The chromatograms were plotted as a percentage relative to the first eluting peaks for each injection. **D**, SDS-PAGE gel separation of the TMPRSS2:nafamostat and TMPRSS2:KD1 complexes. Samples were either heated and reduced (+) or not heated and not reduced (-) prior to gel separation.

### A protein engineering and purification method to isolate the HAI-2 KD2 domain

The KD2 domain has substantial sequence differences between HAI-1 and HAI-2, but the HAI-2 KD2 domain has largely resisted biochemical investigations in the past due to challenges in producing it recombinantly^74^. We designed soluble forms of the complete HAI-2 ectodomain and KD2 domain, but obtained no KD2 protein band in test expression studies (Fig. 6A, and B). To study the isolated KD2 domain in biochemical and biophysical assays, we engineered a cleavable form of the HAI-2 ectodomain to release its KD2 domain after purification (Fig. 6C). We installed a PreScission Protease cut site, naming the construct PreHAI-2. The PreHAI-2 protein was captured from Sf9 insect cell media using Ni^2+^-NTA IMAC and further purified through SEC (Fig. S3A). The PreHAI-2 protein was incubated with PreScission protease to release smaller KD1-His and untagged KD2 protein fragments that were separable by another round of SEC (Fig. S3B). This protein purification scheme ultimately provided pure, untagged forms of the HAI-2 KD1 and KD2 proteins (Fig. 6C).

**Figure 6.**
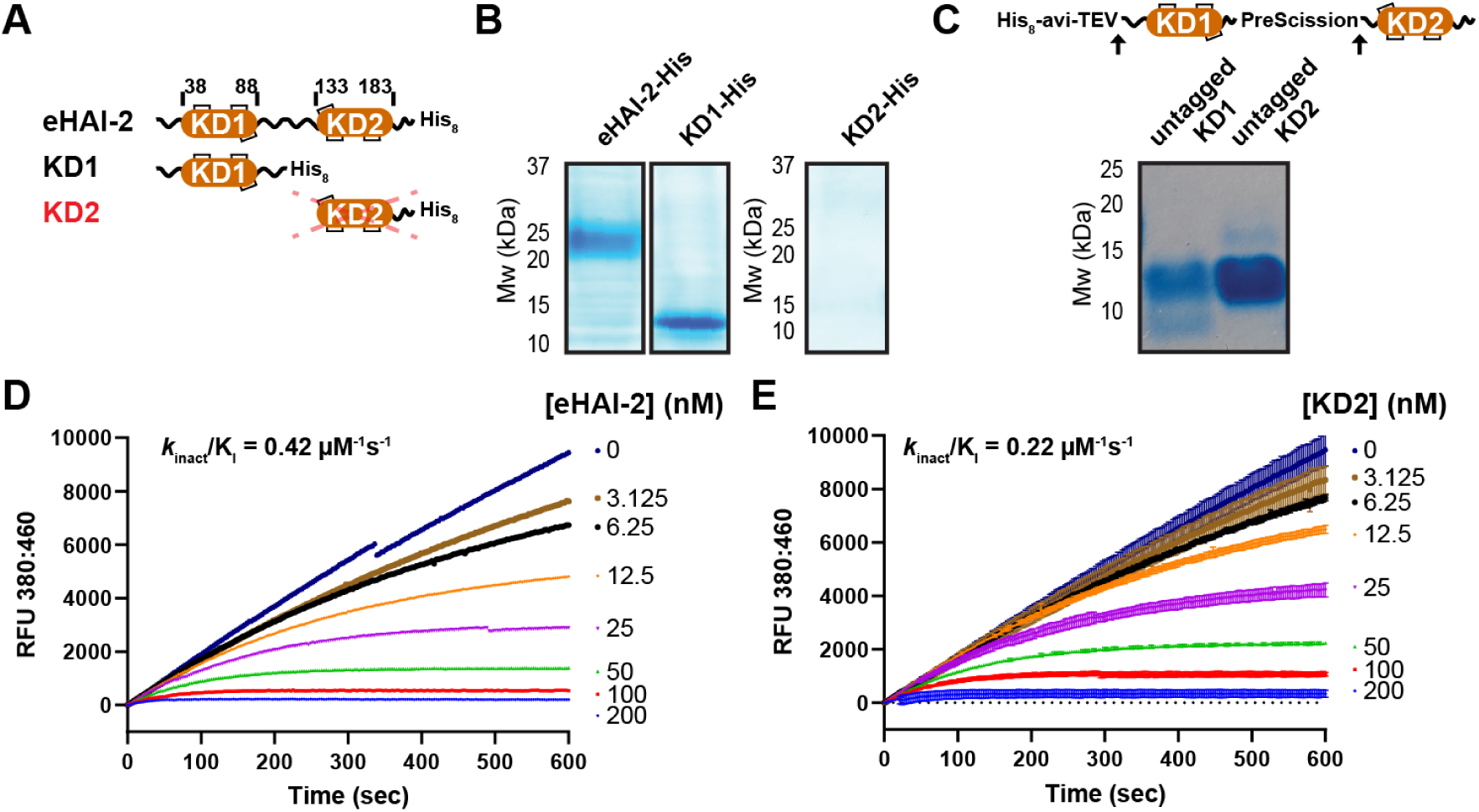
The complete HAI-2 ectodomain and its KD2 domain potently inactivate TMPRSS2 protease activity. **A**, Schematic of recombinant, soluble HAI-2 proteins with C-terminal His tags. **B**, Protein test expression studies for the indicated HAI-2 protein constructs. Secreted protein was purified from 4 mL baculovirus-infected Sf9 insect cells using Ni^2+^-IMAC. **C**, Schematic of an engineered HAI-2 protein with a PreScission protease cut site installed between the KD1 and KD2 domain (PreScission HAI-2; PreHAI-2). Following PreScission cleavage, the released KD1 and KD2 domains (“untagged KD1/KD2”) were purified (below). **D**, Reaction progress curves for TMPRSS2 with the indicated concentrations of eHAI-2 protein or (**E**) KD2 protein added to TMPRSS2 simultaneously with 100 µM Boc-QAR-AMC substrate. Progress curve fitting was performed using the DynaFit software to determine TMPRSS2 *k*_inact_/K_I_ values (Experimental Procedures).

### The HAI-2 KD2 domain rapidly inactivates TMPRSS2

With the untagged eHAI-2, KD1, and KD2 proteins in hand, we compared their TMPRSS2 inhibitory potencies. All three HAI-2 proteins were potent inhibitors of TMPRSS2 peptidase activity, with IC_50_s of (3.0±0.8), (7±3), and (3.9±0.7) nM for eHAI-2, KD1, and KD2, respectively (Fig. S4A). TMPRSS2 reaction progress curves also plateaued in the presence of the eHAI-2 and KD2 proteins, with TMPRSS2 (*k*_inact_/K_I_) values of 0.42 and 0.22 s^-1^µM^-1^, respectively (Fig. 6*D*, *and E*). The eHAI-2 and KD2 proteins induced large TMPRSS2 ΔT_m_s of (27.5±0.4)°C and (24.0±0.1)°C, respectively (Fig. S4B). These data confirm that both the KD1 and KD2 domains of HAI-2 can independently form inactive ternary complexes with TMPRSS2, unlike HAI-1 which can only inhibit TTSPs with its KD1 domain^62^.

### Multivalent HAI-2-IgG fusion proteins are potent TTSP inhibitors

To improve the versatility of HAI-2 proteins as potential anti-TTSP antiviral agents, we developed a series of immunoglobulin G (IgG; Fc)-tagged HAI-2 protein chimeras and expressed them as secreted proteins from HEK293 cells (Expi293F). This HAI-2 protein production strategy had two advantages over our insect-derived HAI-2 proteins; i) These HAI-2 proteins likely contain human post-translational modifications, namely at the two *N*-glycosylation sites at Asn57 and Asn94 (Fig. 7*A*) and ii) disulfide bond formation enables more KDs to be presented per protein molecule (avidity effect).

**Figure 7.**
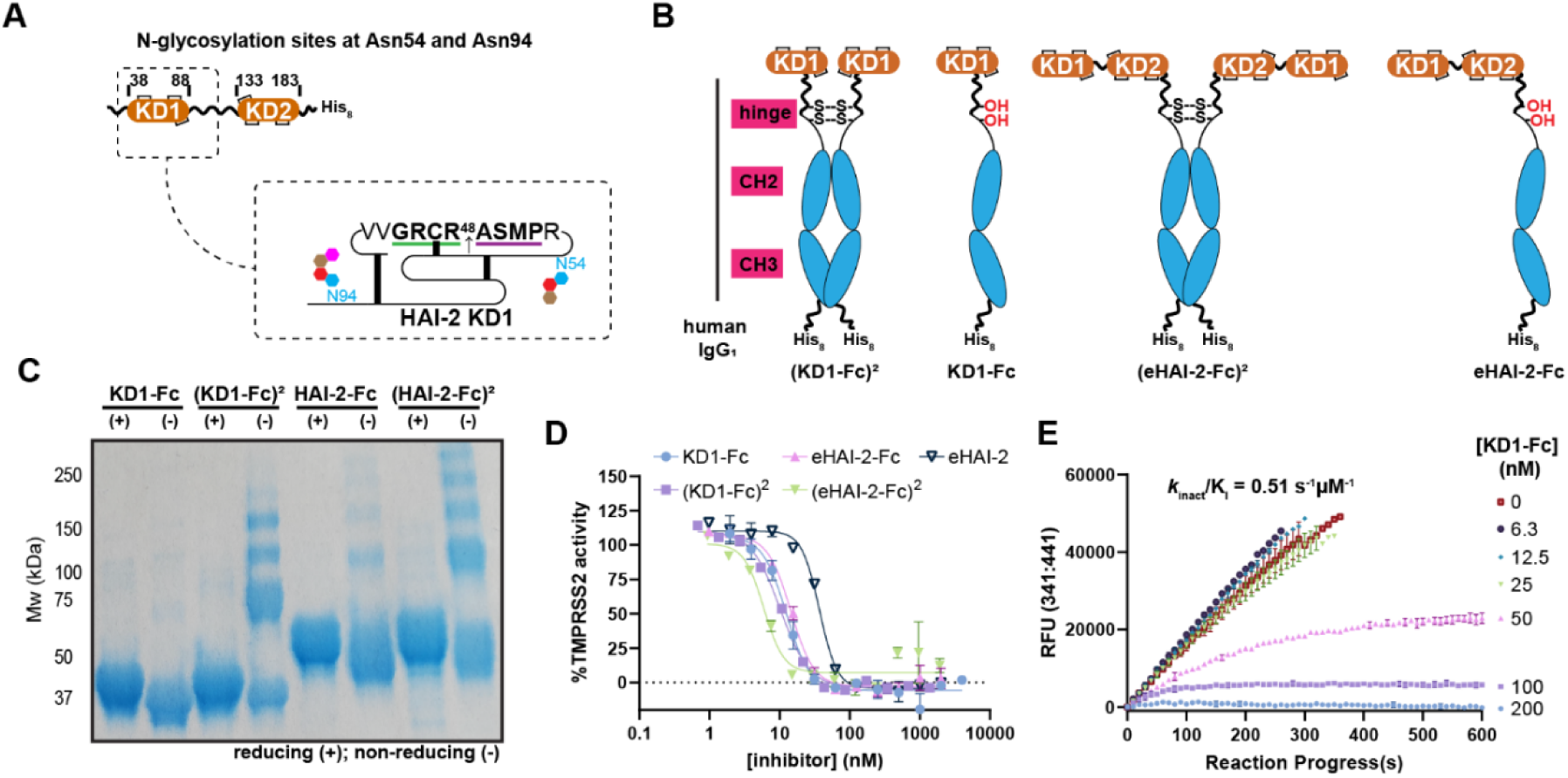
Chimeric HAI-2 proteins produced in HEK293S cells are multivalent TMPRSS2 inhibitors. **A**, Schematic of the N-glycosylation sites present within the HAI-2 protein that are localized around the HAI-2 KD1 domain. A zoomed-in, 2D schematic of the HAI-2 KD1 domain shows the relative locations of the two N-glycosylation sites around the HAI-2 KD1 domain. **B**, Schematics of human Immunoglobulin G (IgG; Fc)-tagged HAI-2 protein chimeras. HAI-2 proteins containing either the isolated KD1 or the HAI-2 ectodomain (eHAI-2) were fused at their C-terminus with Fc tags that were themselves His-tagged. Cys220Ser and Cys226Ser mutations (red text) within the hinge region of the Fc protein were implemented to prevent dimerization. **C**, SDS-PAGE analysis of the indicated HAI-2 protein following secretion from HEK293S cells and Ni^2+^-NTA IMAC purification of HEK293S cell media. The Ni^2+^-NTA IMAC elution samples were thermally denatured and reduced (+) or not thermally denatured and not reduced in advance of SDS-PAGE separation. **D**, TMPRSS2 IC_50_ plots with the indicated HAI-2 proteins. Assays were conducted with 15 nM dasTMPRSS2 protease and 100 µM Boc-QAR-AMC substrate. Initial reaction velocities were tabulated across the first 60 seconds of the assay. The indicated concentrations of inhibitor were pre-incubated with dasTMPRSS2 for 5 minutes prior to the start of the assay. **E**, TMPRSS2 inhibitor-substrate coaddition assay with the KD1-Fc inhibitor and 100 µM Boc-QAR-AMC substrate. Data was curve fit to determine the TMPRSS2 *k*_inact_/K_I_ value using DynaFit 4.0.

To evaluate the effects of having dimeric or monomeric Fc-tagged HAI-2 chimeras containing a single copy of Kunitz Domains or a multivalent set of Kunitz Domains, respectively, we introduced Cys220Ser and Cys226Ser mutations within the hinge region to prevent IgG dimer formation (Fig. 7*B*). Thus, 4 HAI-2 chimeras were designed that contained the monomeric Fc-tagged KD1 domain (KD1-Fc), the dimeric Fc-tagged KD1 domain (KD1-Fc)^2^, the monomeric Fc-tagged eHAI-2 (eHAI-2-Fc), and the dimeric Fc-tagged eHAI-2 ((eHAI-2-Fc)^2^; Fig. 7*B*). The HAI-2 chimeras were overexpressed in HEK293S cells, and the secreted proteins were captured from cell media using Ni^2+^-NTA IMAC purification. The proteins were purified with a single purification step and had exceptionally high purified protein yields for KD1-Fc, (KD1-Fc)^2^, eHAI-2-Fc, and (eHAI-2-Fc)^2^ at 40, 18, 60, and 36 mg per litre HEK293S cell cultures, respectively (Fig. 7*C*). The monomeric HAI-2-Fc chimeras proteins migrated as a single protein band at their expected molecular weight when prepared for SDS-PAGE gel separation using nonreducing and reducing Laemelli buffer. In contrast, the dimeric HAI-2-Fc protein chimeras migrated as several high molecular bands under nonreducing conditions but migrated as a single protein band at their expected molecular weight under reducing conditions (Fig. 7*C*). When the SEC profiles of the monomeric and dimeric KD1-Fc proteins were overlaid, there is a prominent later-eluting peak for the monomeric KD1-Fc protein in both samples, and an earlier peak for the (KD1-Fc)^2^ sample that corresponds to the intact dimer (Fig. S5*A*). When the eHAI-2-Fc and (eHAI-2-Fc)^2^ SEC chromatograms were overlaid, peaks that correspond to the monomeric and dimeric species are evident for the (eHAI-2-Fc)^2^ sample (Fig. S5*B*). The final purified Fc-HAI-2 proteins are shown in Fig. S5*C*. The T_m_s for KD1-Fc, (KD1-Fc)^2^, eHAI-2-Fc, and (eHAI-2-Fc)^2^ were (53.6±0.2), (49.2±0.2), (52.5±0.2), and (48.7±0.3)°C, respectively (Fig. *S5C*).

The TMPRSS2 IC_50_s for KD1-Fc, (KD1-Fc)^2^, eHAI-2-Fc, and (eHAI-2-Fc)^2^ were (12±3), (9.9±0.9), (14±1.5), and (6±2) nM, respectively (Fig. 7*D*). In comparison, the eHAI-2 protein produced from baculovirus-infected insect cells had a TMPRSS2 IC_50_ of (36±3) nM when run in the same assay. Furthermore, the TMPRSS2 *k*_inact_/K_I_ value for KD1-Fc was 0.51 s^-1^µM^-1^ (Fig. 7*E*). These data confirm that Fc-tagged HAI-2 proteins can be produced in Expi293F cells and additional KD1 or KD2 domains can enhance the binding affinity (through avidity) toward TMPRSS2.

### Soluble HAI-2 proteins broadly inactivate airway TTSPs but not thrombin

To evaluate if HAI-2 proteins broadly and potently disable airway TTSP activity, we screened TMPRSS2/3/13/11A/11B/11D/11E/11F activity in the presence of KD1-Fc. The Fc-tagged KD1 protein was tested at a concentration of 100 nM, resulting in potent inhibition for all tested proteases except TMPRSS11B and TMPRSS11E (Fig. 8*A*). Within the same assay, all TTSPs were inhibited by a control inhibitor 6-amidino-2-naphthol (50 µM). Interestingly, trypsin, but not thrombin, was inhibited by the HAI-2 protein, confirming that only some trypsin-like serine proteases are susceptible to inhibition by HAI-2. We confirmed these findings in an orthogonal assay using SARS-CoV-2 S protein as a substrate for the protease activities of TMPRSS2, TMPRSS15, thrombin, and trypsin (Fig. 8*B*). In the absence of protease inhibitor, TMPRSS2 and thrombin cleaved the S protein and eliminated the intact S protein band. Nafamostat, KD1, and KD2 completely blocked TMPRSS2 and trypsin activity towards the S protein. In contrast, thrombin was inhibited by nafamostat and not KD1 or KD2. These data collectively indicated that airway TTSPs could be targeted with HAI-2 proteins, and HAI-2 proteins may avoid inhibiting other critical trypsin-like serine proteases found in human blood.

**Figure 8.**
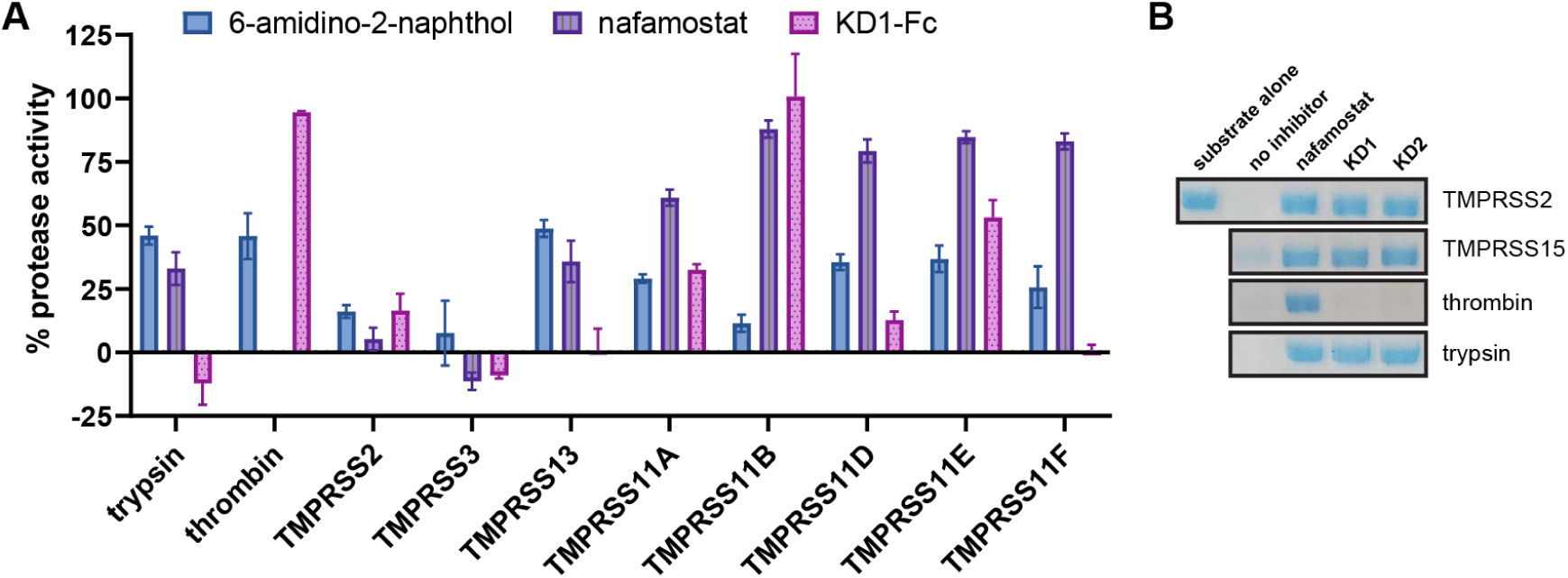
The HAI-2 KD1 domain broadly disables many TTSPs but not thrombin. **A**, Protease activity (relative to vector control) in the presence of 6-amidino-2-naphthol (50 µM), nafamostat (50 nM), or KD1-Fc (100 nM). Assays contained 100 µM Boc-QAR-AMC substrate. Substrate and inhibitor were transferred simultaneously and reaction progress monitored for 15 minutes. **B**, SARS-CoV-2 S protein cleavage assay with 25 nM dasTMPRSS2, 80 Units (U) bovine TMPRSS15 (NEB Cat #P8070L), 0.5 U human thrombin (Millipore Cat # 605195-5000U), and 420 nM bovine pancreatic trypsin (Sigma Cat # T8003-100MG). The proteases were preincubated with 2 µM inhibitor for 10 min prior to transfer to 4 µg S protein substrate. S protein was incubated with dasTMPRSS2 protease at 20°C for 10 minutes. Samples were prepared for SDS-PAGE through addition of 4X nonreducing Laemelli buffer and thermally denatured (95°C for 5 minutes), then separated on a 4-20% Tris-Glycine SDS-PAGE gel and stained with Coomassie blue.

## Discussion

Several cell-based studies investigating HAI-2 as a regulator of TMPRSS2 protease activity in the context of cancer and viral infections, but translation efforts with soluble HAI-2-based therapeutic strategies have not been explored yet. In this mechanism-focused study, we interrogated the unique strategy by which HAI-2 (and probably HAI-1) inactivate TMPRSS2 and other TTSPs by exclusively forming a stable ternary complex with the activated protease, but the P1-P1’ peptide bond is not nucleophilically attacked by the catalytic serine residue. This is different from other macromolecular inhibitors of trypsin-like serine proteases that employ the Standard inhibition mechanism, including BPTI^68,69^, SFTI-1^66,75–77^, and SPINK1/2^70,71^ which have reactive peptide bonds to trap their target proteases^65^. The HAI-2 protein has broad activity against most TTSPs but avoided inhibition of other trypsin-like serine proteases like thrombin. This is an encouraging property for the translation of these proteins as an anti-TTSP drug strategy. Protein-based inhibitors of serine proteases have been explored as therapeutics in numerous clinical trials for respiratory disorders, including chronic obstructive pulmonary disorder (COPD)^78^, alpha-1 antitrypsin (AAT) deficiency disorder^78^, cystic fibrosis^79–81^, and COVID-19^82^. In several clinical trials, recombinant serine protease inhibitors were administered as aerosols by nebulizing a solution of the protein drug^78–82^. Accordingly, there are delivery mechanisms under development for administering therapeutic concentrations of macromolecular serine protease inhibitors to the human airways^83^ that could potentially be reapplied for HAI-2.

A previous study proposed that HAI-2 binds to TMPRSS2 prior to zymogen activation, preventing TMPRSS2 zymogen activation and in effect blocked cell surface TMPRSS2 activity^61^. To measure the biophysical interaction between the zymogen form of TMPRSS2 with the HAI-2 protein, we developed a “zymogen-locked” form of TMPRSS2 (TMPRSS2 DDDDA255). This was necessary because we have observed partial zymogen cleavage activation for TMPRSS2 R255Q and TMPRSS2 S441A (unpublished data) which have previously been reported to prevent zymogen activation^15,84^. There are a few alternative hypotheses that do not require HAI-2 binding to zymogen TMPRSS2 but explain why active TMPRSS2 protein levels may be decreased upon induction of HAI-2 protein expression and are consistent with the biophysical data presented in this work. Firstly, increased HAI-2 protein levels could block the activity of the active protease(s) driving TMPRSS2 zymogen activation. We have recently shown that TMPRSS2 can autoactivate^43^, so increased HAI-2 protein levels may block an initiator TMPRSS2 molecule from activating other TMPRSS2 zymogen molecules^43^. Alternatively, active TMPRSS2 may have rapidly undergone zymogen activation, formed a complex with the excess HAI-2, and the TMPRSS2:HAI-2 complex was shed from the cell surface. This shedding mechanism was examined in detail for matriptase with HAI-1, where ectopic HAI-1 overexpression reduced matriptase levels at the cell surface and increased the levels of the matriptase:HAI-1 complex in cell media^24^. Finally, overexpression of HAI-2 and reduction of active TMPRSS2 protein levels (detected through western blotting) could be explained by complex formation with HAI-2, masking the TMPRSS2 epitope recognized by a TMPRSS2-specific primary antibody. We demonstrated that heat treatment was required to dissociate TMPRSS2:HAI-2 complexes in advance of SDS-PAGE gel separation and detect the active TMPRSS2 protein band (Fig. *5D*). Thus, while the Zhang *et al.* (2022) study suggested HAI-2 prevents TMPRSS2 activation by binding before zymogen activation, alternative explanations such as inhibition of the TMPRSS2-activating protease, rapid TMPRSS2:HAI-2 complex formation and shedding, or epitope masking due to complex formation are also plausible and align with the data presented here.

The TMPRSS2-HAI-2 inhibition kinetics and biophysical binding data confirmed that the Kunitz Domains of HAI-2 disable active TMPRSS2 by forming inactive ternary complexes that do not require the TMPRSS2 S441 residue. Our TMPRSS2 coaddition assays with HAI-2 proteins and substrate allowed us to determine the *k*_inact_/K_I_ values for TMPRSS2 inactivation through reaction progress curve fitting. A previous publication showed that matriptase reaction progress curves plateaued when truncated HAI-1 proteins were briefly incubated with matriptase prior to addition of substrate, suggesting that HAI-1 may inactivate matriptase using a similar inhibition mechanism^36^. By introducing mutations to the HAI-2 KD1 residue A49 (GRCR48<u>A</u>SMP), we modified a key residue involved in the P1-P1’ peptide bond and perturbed the TMPRSS2 inhibition mechanism. Two HAI-2 KD1 A49X mutants, KD1 A49G and KD1 A49E, served as useful surrogates to also understand the TMPRSS2 inhibition kinetics with the HAI-1 KD1 and HAI-1 KD2 domains, respectively. The HAI-2 KD1 A49G protein potently inhibited the activity of TMPRSS2, matching previous HAI-1 KD1 (GRCR260GSFP) inhibition data reported for TMPRSS2^46^, matriptase^48,49,53,85^, and prostasin^48,49,53,85,86^. In contrast, the HAI-2 KD1 A49E protein did not inhibit TMPRSS2 activity, which is consistent with the finding that the HAI-1 KD2 domain (GLCK401ESIP) does not inhibit the protease activity of TTSPs^36^.

Unlike HAI-1, HAI-2 is capable of inhibiting TTSPs using either its KD1 or KD2 domain. The biological implications of this two-pronged inhibition feature are unclear. The HAI-2 KD1 and KD2 domains are connected by a long linker region that is expected to be unstructured, which may allow the distance and flexibility to allow for each KD to engage a separate protease molecule at the cell surface. Our SDS-PAGE analysis of the TMPRSS2:KD1 complex confirmed that it was stable to mild SDS treatment in advance of gel separation, but the complex dissociated if the sample was heated. Thus, western blotting for these complexes require specific sample preparation to increase the likelihood of detection.

The HAI-2 KD1 A49M and A49T proteins blocked TMPRSS2 activity but dissociated over time. It remains unclear whether a permanent inhibitor or competitive inhibitor would be the most effective for blocking TMPRSS2 activity in antiviral treatments^87–89^. Thus, the identification of these engineered KD1 mutants provides a potentially useful and an alternative strategy to traditional small molecule or peptide-based molecules that disable TMPRSS2 through competitive inhibition, reversible covalent inhibition (ketobenzothiazole-bearing peptidomimetics), or permanent covalent inhibition (chloromethylketone-bearing peptidomimetics). To confer greater TMPRSS2 selectivity, additional mutations to the KD1 domain will be required to distinguish TMPRSS2 over other TTSPs. However, several TTSPs have been shown to drive SARS-CoV-2 infection in human airway cell lines, including TMPRSS11A and TMPRSS11D^9,11,12,32,39,90^. There may be multiple TTSPs that collaborate or are functionally redundant for driving respiratory virus infections in humans. Thus, a pharmacological, pan-TTSP intervention strategy may provide more protective effects for human respiratory virus infections than a drug targeting a single TTSP. Future chemical biology studies that dissect the function of each TTSP may inform a smaller list of critical TTSPs that are essential for infection and thereby focus drug development efforts.

## Materials and Methods

### Construct design and cloning

HAI-2 cDNA (nucleotide accession # BC125196) was accessed from the Mammalian Gene Collection. The entire HAI-2 ectodomain (residues 29-197; eHAI-2) was subcloned into the pFHMSP-LIC C baculovirus donor vector that encoded a N-terminal honeybee melittin signal sequence peptide and C-terminal His_8_-tag with mutagenesis primers available in Extended Data Table 1. For Fc-tagged HAI-2 protein constructs designed for protein production in Expi293F cells, constructs were subcloned into a pcDNA3 mammalian expression vector containing a N-terminal MRPTWAWWLFLVLLLALWAPARG artificial secretion signal peptide^91^, as well as a C-terminal, human IgG (Fc) protein and His_8_ tag. To disrupt dimerization through Fc-mediated disulfide bond formation (“mono-Fc”), Cys220Ser and Cys226Ser mutations were introduced to the Fc protein.

For HAI-2 protein produced in Sf9 cells using the baculovirus expression vector system (BEVS), plasmid transfer vector containing *HAI-2* was initially transformed into *Escherichia coli* DH10Bac cells (Thermo Fisher; Cat# 10361012) to generate recombinant viral bacmid DNA. Sf9 cells were transfected with Bacmid DNA using JetPrime transfection reagents (PolyPlus Transfection Inc.; Cat# 114-01) according to the manufacturer’s instructions, and recombinant baculovirus particles were obtained and amplified from P1 to P2 viral stocks. Recombinant P2 viruses were used to generate suspension culture of baculovirus infected insect cells (SCBIIC) for scaled-up production of HAI-2 proteins.

For HAI-2 proteins produced in Expi293F cells, FectoPro transfection reagent (Polyplus-transfection SA, Cat. #116-010) was used to transfect plasmid DNA to >95% viable cells in the presence of sodium butyrate according to the manufacturer’s instructions.

### HAI-2 protein production in Sf9 and Expi293F cells

Sf9 cells were grown in I-Max Insect Medium (Wisent Biocenter; Cat# 301-045-LL) to a density of 4x10^6^ cells/mL and infected with 20 mL/L of suspension culture of baculovirus infected insect cells prior to incubation on an orbital shaker (145 rpm, 26 °C). Expi293F cells were grown in Expi293 Expression Medium (Life Techn. Cat.# A1435102) for approximately 72 hours post-transfection.

### Recombinant HAI-2 protein purifications

HAI-2 proteins were produced through secreted expression and purified using similar protocol that was previously described for TMPRSS2. Cell culture medium containing the final secreted protein products AA-[HAI-2]-EFVEHHHHHHH were collected by centrifugation (20 min, 10 °C, 6,000 x g) 4-5 days post-infection when Sf9 cell viability dropped to 55 - 60%. For Expi293F-produced HAI-2 proteins, cell culture medium was collected approximately 72 hours post-transfection. Media was adjusted to pH 7.4 by addition of concentrated PBS stock, then supplemented with 15 mL/L settled Ni-NTA resin (Qiagen) at a scale of 1-4 L. A single batch Ni^2+^-NTA IMAC purification were used to capture protein, with each round requiring shaking in 2.8L flasks for 1 hour at 16 °C (110 rpm), then transferred to a 1.0 L gravity flow column (Bio-Rad). Beads were washed with 3 column volumes (CVs) PBS prior to elution with 1.5 resin bed volumes of Elution Buffer (PBS supplemented with 250 mM imidazole). Crude protein was concentrated using 10 kDa MWCO Amicon filters for the eHAI-2 and Fc-HAI-2 proteins or 3 kDa MWCO Amicon filters for KD1 proteins. Protein samples were prepared for SDS-PAGE with either reducing (5 mM 2-mercaptoethanol) Laemelli dye and thermally denatured for 5 minutes, or nonreducing Laemelli dye not subjected to thermal denaturation. Concentrated Ni^2+^-NTA IMAC elution samples were passed through a 0.22 µm syringe filter and injected to a Superdex75 gel filtration column pre-equilibrated with Size-Exclusion Chromatography (SEC) Buffer (50 mM Tris pH 8.0, 200 mM NaCl). SEC fractions containing HAI-2 proteins were pooled, concentrated to 5 mg/mL, then flash frozen in liquid nitrogen and stored as aliquots at -80°C in advance of enzyme assays.

### Protein sequence alignments and structure superpositions

The FASTA sequences of human HAI-1 and HAI-2 proteins were accessed through Uniprot (isoform 1) and aligned using Clustal Omega and annotated with ESPript v.3.0. Protein structures were also accessed from the PDB and superposed using PyMOL Protein structure figures were prepared in PyMOL.

### Enzyme peptidase assays and IC_50_ determination

Peptidase inhibition assays with fluorogenic Boc-Gln-Ala-Arg-AMC substrate (Bachem Cat # 4017019.0025) were carried out as described previously for dasTMPRSS2^72^. Assays contained a final concentration of 6 nM dasTMPRSS2 with buffer containing 50 mM Tris pH 8.0 and 200 mM NaCl. Nafamostat mesylate (MedChemExpress Cat# HY-B0190A) and 6-amidino-2-naphthol methanesulfonate (TCI Cat# A1193) were prepared as fresh DMSO stocks immediately prior to inhibition assays. To determine half-maximal inhibitory concentrations (IC_50_s), inhibitor titrations were prepared in DMSO for nafamostat and 6-amidino-2-naphthol or SEC buffer for HAI-2 proteins. Inhibitor stocks were transferred to wells containing dasTMPRSS2 enzyme and incubated for 10 min. Boc-QAR-AMC substrate was transferred to wells (100 µM final concentration) and plates immediately read for fluorescence. Initial reaction velocity slopes were tabulated across the first 0-120 seconds of the assay and normalized relative to uninhibited enzyme. Curves were fit for Absolute IC_50_ in Graphpad Prism.

### TMPRSS2 *k*_inact_/K_I_ and K_i_ kinetic assays and curve fitting

The TMPRSS2 inactivation rate (*k*_inact_/K_I_) with nafamostat and HAI-2 inhibitors was measured using protocols and Dynafit4 curve fitting scripts described for TMPRSS11D^43^. Stocks of Boc-QAR-AMC substrate with various concentrations of nafamostat, 6-amidino-2-naphthol, or HAI-2 proteins were transferred to wells containing dasTMPRSS2 enzyme and fluorescence immediately read to capture the rapid acylation. Wells contained final concentrations of 100 µM Boc-QAR-AMC substrate and 6 nM dasTMPRSS2. Reaction curve traces and DynaFit enzyme kinetic scripts are available in Supplementary Figures 3-4.

### SARS-CoV-2 S protein cleavage inhibition assays

The SARS-CoV-2 S protein construct HexaFurin was prepared as a substrate for dasTMPRSS2 and commercially available serine proteases using a similar protocol to that described for dasTMPRSS11D^43^. Inhibitors were pre-incubated with proteases for 10 min prior to transferring the enzyme-inhibitor mixtures to 5 µg S protein substrate in Tris pH 8.0 with 200 mM NaCl and final protease concentrations of 0.01 mg/mL for trypsin (Sigma Cat # T8003-100MG), chymotrypsin, 200 nM for dasTMPRSS2, and 80 U enteropeptidase (NEB Cat #P8070L), and 0.5 U thrombin. S protein digestions took place in microplates submerged in a 37 °C water bath for 1 hr before the assay was terminated through addition of 100 μM nafamostat. Samples were prepared for SDS-PAGE through addition of Laemmeli sample buffer and were thermally denatured at 95 °C for 5 min prior to gel separation. Gels stained with Coomassie blue prior to destaining and visualization.

### DSF for ligand-induced melting temperature shifts

TMPRSS2 protein melting temperatures (T_m_s) and ligand-induced T_m_ shifts were measured using SYPRO Orange dye (Life Technologies, catalog no. S-6650) and monitoring fluorescence at 470 and 510 nm excitation and emission, respectively, using the Light Cycler 480 II (Roche Applied Science). Samples were prepared in at least technical triplicate in 384-well plates (Axygen; catalog nos. PCR-384-C; UC500) at a final volume of 20 µL. Experiments contained 2.5 µM TMPRSS2 protein or 2 µM TMPRSS2 S441A protein, 10% (*v*/*v*) ligand (or vector control) and 5× SYPRO Orange. Thermal melt curves were generated across a 25 to 95 °C gradient at a rate of 2 °C/min. Protein melt curves were normalized to the maximum observed fluorescence and plotted in Graphpad Prism. Protein T_m_s were calculated using the dRFU method with the DSFworld application^92^. Ligand-induced TMPRSS2 or TMPRSS2 S441A shifts (ΔT_m_s) relative to vector control were calculated for each concentration of ligand and plotted using Graphpad Prism.

## Author Contributions

C.H.A., B.J.F., G.B.M., and F.B. provided project supervision; B.J.F conceived the project; B.J.F. and R.P.W. designed the experiments; Y.L. cloned protein constructs for expression; P.L. generated bacmids; A.S., R.P.W., O.I., A.H., Y-Y.L., and M.K. produced proteins in insect cells; B.J.F., O.I., and J.L. purified recombinant proteins; B.J.F. performed bioinformatic and structural analyses; B.J.F., R.P.W. and J.L. performed fluorogenic peptidase activity and inhibitor potency assays and analyzed kinetics; B.J.F. and O.I. performed gel-based S protein digestion assays; B.J.F., O.I., and J.L. performed DSF assays; B.J.F. and R.P.W. prepared figures; B.J.F, R.P.W. and C.H.A wrote the manuscript.

## Acknowledgments

This work was supported by the BC Leadership Chair in Functional Cancer Imaging to F.B., the Canadian Institute of Health Research grant (no. FDN154328) to C.H.A., and a Mitacs Elevate Postdoctoral Fellowship to B.J.F. The Structural Genomics Consortium is a registered charity (no. 1097737) that receives funds from Bayer AG, Boehringer Ingelheim, Bristol Myers Squibb, Genentech, Genome Canada through Ontario Genomics Institute (grant no. OGI-196), the EU and EFPIA through the Innovative Medicines Initiative 2 Joint Undertaking (EUbOPEN grant no. 875510), Janssen, Merck KGaA (also known as EMD in Canada and the United States), Pfizer and Takeda.

## References

1. Du, X. et al. The serine protease TMPRSS6 is required to sense iron deficiency. Science (1979). 320, 1088–1092 (2008).

2. List, K. et al. Epithelial integrity is maintained by a matriptase-dependent proteolytic pathway. American Journal of Pathology 175, 1453–1463 (2009).

3. List, K. et al. Autosomal ichthyosis with hypotrichosis syndrome displays low matriptase proteolytic activity and is phenocopied in ST14 hypomorphic mice. Journal of Biological Chemistry 282, 36714–36723 (2007).

4. Limburg, H. et al. TMPRSS2 is the major activating protease of Influenza A virus in primary human airway cells and influenza B virus in human type II pneumocytes. J. Virol. 93, 1–22 (2019).

5. Böttcher, E. et al. Proteolytic Activation of Influenza Viruses by Serine Proteases TMPRSS2 and HAT from Human Airway Epithelium. J. Virol. 80, 9896–9898 (2006).

6. Okumura, Y. et al. Novel Type II Transmembrane Serine Proteases, MSPL and TMPRSS13, Proteolytically Activate Membrane Fusion Activity of the Hemagglutinin of Highly Pathogenic Avian Influenza Viruses and Induce Their Multicycle Replication. J. Virol. 84, 5089–5096 (2010).

7. Mahoney, M., et al. A novel class of TMPRSS2 inhibitors potently block SARS-CoV-2 and MERS-CoV viral entry and protect human epithelial lung cells. PNAS 118, e2108728118 (2021).

8. Reinke, L. M. et al. Different residues in the SARS-CoV spike protein determine cleavage and activation by the host cell protease TMPRSS2. PLoS One 12, 1–15 (2017).

9. Kishimoto, M. et al. TMPRSS11D and TMPRSS13 activate the SARS-CoV-2 spike protein. Viruses 13, 384 (2021).

10. Iwata-Yoshikawa, N. et al. Essential role of TMPRSS2 in SARS-CoV-2 infection in murine airways. Nat. Commun. 13, 6100 (2022).

11. Zang, R. et al. TMPRSS2 and TMPRSS4 promote SARS-CoV-2 infection of human small intestinal enterocytes. Sci. Immunol. 5, 1–14 (2020).

12. Bestle, D. et al. TMPRSS2 and furin are both essential for proteolytic activation of SARS-CoV-2 in human airway cells. Life Sci Alliance 3, e202000786 (2020).

13. Sasaki, M. et al. SARS-CoV-2 variants with mutations at the S1 / S2 cleavage site are generated in vitro during propagation in TMPRSS2-deficient cells. PLoS Pathog. 17, e1009233 (2021).

14. Klezovitch, O. et al. Hepsin promotes prostate cancer progression and metastasis. Cancer Cell 6, 185–195 (2004).

15. Chen, Y. et al. TMPRSS2, a Serine Protease Expressed in the Prostate on the Apical Surface of Luminal Epithelial Cells and Released into Semen in Prostasomes, Is Misregulated in Prostate Cancer Cells. American Journal of Pathology 176, 2986–2996 (2010).

16. Ko, C. J. et al. Androgen-induced TMPRSS2 activates matriptase and promotes extracellular matrix degradation, prostate cancer cell invasion, tumor growth, and metastasis. Cancer Res. 75, 2949–2960 (2015).

17. Herter, S. et al. Hepatocyte growth factor is a preferred in vitro substrate for human hepsin, a membrane-anchored serine protease implicated in prostate and ovarian cancers. Biochemical Journal 390, 125–136 (2005).

18. Larzabal, L. et al. Overexpression of TMPRSS4 in non-small cell lung cancer is associated with poor prognosis in patients with squamous histology. Br. J. Cancer 105, 1608–1614 (2011).

19. Chikaishi, Y. et al. TMPRSS4 Expression as a Marker of Recurrence in Patients with Lung Cancer. Anticancer Res. 36, 121–127 (2016).

20. Tsai, C. H. et al. HAI-2 suppresses the invasive growth and metastasis of prostate cancer through regulation of matriptase. Oncogene 33, 4643–4652 (2014).

21. Mukai, S. et al. Matriptase and MET are prominently expressed at the site of bone metastasis in renal cell carcinoma: immunohistochemical analysis. Hum. Cell 28, 44–50 (2015).

22. Wang, J. Y., Jin, X. & Li, X. F. Knockdown of TMPRSS3, a transmembrane serine protease, inhibits proliferation, migration, and invasion in human nasopharyngeal carcinoma cells. Oncol. Res. 26, 95–101 (2018).

23. Bugge, T. H., Antalis, T. M. & Wu, Q. Type II transmembrane serine proteases. Journal of Biological Chemistry 284, 23177–23181 (2009).

24. Tseng, C. et al. Matriptase shedding is closely coupled with matriptase zymogen activation and requires de novo proteolytic cleavage likely involving its own activity. PLoS One 12, 1–23 (2017).

25. Lee, M. S. et al. Autoactivation of matriptase in vitro: Requirement for biomembrane and LDL receptor domain. Am. J. Physiol. Cell Physiol. 293, 95–105 (2007).

26. Knappe, S., Wu, F., Masikat, M. R., Morser, J. & Wu, Q. Functional analysis of the transmembrane domain and activation cleavage of human corin: Design and characterization of a soluble corin. Journal of Biological Chemistry 278, 52363–52370 (2003).

27. Wang, H. et al. Distinct roles of N-glycosylation at different sites of corin in cell membrane targeting and ectodomain shedding. Journal of Biological Chemistry 290, 1654–1663 (2015).

28. Ohno, A. et al. Crystal structure of inhibitor-bound human MSPL that can activate high pathogenic avian influenza. Life Sci. Alliance 4, 1–11 (2021).

29. Tang, T. et al. Proteolytic Activation of SARS-CoV-2 Spike at the S1/S2 Boundary: Potential Role of Proteases beyond Furin. ACS Infect. Dis. 7, 264–272 (2021).

30. Hoffmann, M. et al. SARS-CoV-2 Cell Entry Depends on ACE2 and TMPRSS2 and Is Blocked by a Clinically Proven Protease Inhibitor. Cell 181, 271–280 (2020).

31. Hoffmann, M., Kleine-Weber, H. & Pohlmann, S. A Multibasic Cleavage Site in the Spike Protein of SARS-CoV-2 Is Essential for Infection of Human Lung Cells. Mol. Cell 78, 779–784 (2020).

32. Wruck, W. & Adjaye, J. SARS-CoV-2 receptor ACE2 is co-expressed with genes related to transmembrane serine proteases, viral entry, immunity and cellular stress. Sci. Rep. 10, 1–14 (2020).

33. Shia, S. et al. Conformational lability in serine protease active sites: Structures of hepatocyte growth factor activator (HGFA) alone and with the inhibitory domain from HGFA inhibitor-1B. J. Mol. Biol. 346, 1335–1349 (2005).

34. Zhao, B. et al. Crystal structures of matriptase in complex with its inhibitor hepatocyte growth factor activator inhibitor-1. Journal of Biological Chemistry 288, 11155–11164 (2013).

35. Yamasaki, Y., Satomi, S., Murai, N., Tsuzuki, S. & Fushiki, T. Inhibition of membrane-type serine protease 1/matriptase by natural and synthetic protease inhibitors. J. Nutr. Sci. Vitaminol. (Tokyo). 49, 27–32 (2003).

36. Kojima, K., Tsuzuki, S., Fushiki, T. & Inouye, K. Roles of functional and structural domains of hepatocyte growth factor activator inhibitor type 1 in the inhibition of matriptase. Journal of Biological Chemistry 283, 2478–2487 (2008).

37. Wettstein, L. et al. Alpha-1 antitrypsin inhibits TMPRSS2 protease activity and SARS-CoV-2 infection. Nat. Commun. 12, 1–10 (2021).

38. Jiang, J. et al. Ectodomain shedding and autocleavage of the cardiac membrane protease corin. Journal of Biological Chemistry 286, 10066–10072 (2011).

39. Matsuyama, S. et al. Enhanced isolation of SARS-CoV-2 by TMPRSS2-expressing cells. Proc. Natl. Acad. Sci. U. S. A. 117, 7001–7003 (2020).

40. Heurich, A. et al. TMPRSS2 and ADAM17 Cleave ACE2 Differentially and Only Proteolysis by TMPRSS2 Augments Entry Driven by the Severe Acute Respiratory Syndrome Coronavirus Spike Protein. J. Virol. 88, 1293–1307 (2014).

41. Matsuyama, S. et al. Efficient activation of the severe acute respiratory syndrome coronavirus spike protein by the transmembrane protease TMPRSS2. J. Virol. 84, 12658–12664 (2010).

42. Glowacka, I. et al. Evidence that TMPRSS2 Activates the Severe Acute Respiratory Syndrome Coronavirus Spike Protein for Membrane Fusion and Reduces Viral Control by the Humoral Immune Response. J. Virol. 85, 4122–4134 (2011).

43. Fraser, B. J. et al. Structural basis of TMPRSS11D specificity and autocleavage activation. bioRxiv 10.1101/2024.10.09.617371 (2024) doi:10.1101/2024.10.09.617371.

44. Hoffmann, M. et al. Camostat mesylate inhibits SARS-CoV-2 activation by TMPRSS2-related proteases and its metabolite GBPA exerts antiviral activity. EBioMedicine 65, (2021).

45. Hoffmann, M. et al. Nafamostat mesylate blocks activation of SARS-CoV-2: New treatment option for COVID-19. Antimicrob. Agents Chemother. 64, (2020).

46. Ko, C. et al. Inhibition of TMPRSS2 by HAI-2 reduces prostate cancer cell invasion and metastasis. Oncogene 39, 5950–5963 (2020).

47. Hamilton, B. S. et al. Inhibition of influenza virus infection and hemagglutinin cleavage by the protease inhibitor HAI-2. Biochem. Biophys. Res. Commun. 450, 1070–1075 (2014).

48. Lai, C. H. et al. Matriptase complexes and prostasin complexes with HAI-1 and HAI-2 in human milk: Significant proteolysis in lactation. PLoS One 11, (2016).

49. Lee, S. P. et al. Tissue distribution and subcellular localizations determine in vivo functional relationship among prostasin, matriptase, HAI-1, and HAI-2 in human skin. PLoS One 13, e0192632 (2018).

50. Tomita, Y. et al. The Physiological TMPRSS2 Inhibitor HAI-2 Alleviates SARS-CoV-2 Infection. J. Virol. 95, 11–13 (2021).

51. Ko, C. et al. Inhibition of TMPRSS2 by HAI-2 reduces prostate cancer cell invasion and metastasis. Oncogene 39, 5950–5963 (2020).

52. Rau, J. C., Beaulieu, L. M., Huntington, J. A. & Church, F. C. Serpins in thrombosis, hemostasis and fibrinolysis. Journal of Thrombosis and Haemostasis 5, 102–115 (2007).

53. Holt-Danborg, L. et al. SPINT2 (HAI-2) missense variants identified in congenital sodium diarrhea/tufting enteropathy affect the ability of HAI-2 to inhibit prostasin but not matriptase. Hum. Mol. Genet. 28, 828–841 (2019).

54. Bou Chaaya, S., Eason, J. D. & Ofoegbu, B. N. Syndromic congenital diarrhoea: New SPINT2 mutation identified in the UAE. BMJ Case Rep. 2017, (2017).

55. Heinz-Erian, P. et al. Mutations in SPINT2 cause a syndromic form of congenital sodium diarrhea. Am. J. Hum. Genet. 84, 188–196 (2008).

56. Bergum, C. & List, K. Loss of the matriptase inhibitor HAI-2 during prostate cancer progression. Prostate 70, 1422–1428 (2010).

57. Kongkham, P. N. et al. An epigenetic genome-wide screen identifies SPINT2 as a novel tumor suppressor gene in pediatric medulloblastoma. Cancer Res. 68, 9945–9953 (2008).

58. Pereira, M. S. et al. Loss of SPINT2 expression frequently occurs in glioma, leading to increased growth and invasion via MMP2. Cellular Oncology 43, 107–121 (2020).

59. Yamauchi, M. et al. Expression of hepatocyte growth factor activator inhibitor type 2 (HAI-2) in human testis: identification of a distinct transcription start site for the HAI-2 gene in testis. Biol. Chem. 383, 1953–1957 (2002).

60. Murray, A. S. et al. Phosphorylation of the type II transmembrane serine protease, TMPRSS13, in hepatocyte growth factor activator inhibitor-1 and -2-mediated cell-surface localization. Journal of Biological Chemistry 292, 14867–14884 (2017).

61. Zhang, Y. et al. Transmembrane serine protease TMPRSS2 implicated in SARS-CoV-2 infection is autoactivated intracellularly and requires N-glycosylation for regulation. Journal of Biological Chemistry 298, 102643 (2022).

62. Kirchhofer, D. et al. Tissue expression, protease specificity, and Kunitz domain functions of hepatocyte growth factor activator inhibitor-1B (HAI-1B), a new splice variant of HAI-1. Journal of Biological Chemistry 278, 36341–36349 (2003).

63. Fraser, B. et al. Large Library Docking and Biophysical Analysis of Small-Molecule TMPRSS2 Inhibitors. J. Med. Chem. (2025).

64. Finkenstadt, W. & Laskowski, M. Peptide Bond Cleavage on Trypsin-Trypsin Inhibitor Complex Formation. Journal of Biological Chemistry 240, 962–963 (1965).

65. Laskowski, M. & Kato, I. PROTEIN INHIBITORS OF PROTEINASES. Ann. Rev. Biochem 49, 9–626 (2025).

66. De Veer, S. J. et al. Engineered protease inhibitors based on sunflower trypsin inhibitor-1 (SFTI-1) provide insights into the role of sequence and conformation in Laskowski mechanism inhibition. Biochemical Journal 469, 243–253 (2015).

67. Riley, B. T. et al. KLK4 Inhibition by Cyclic and Acyclic Peptides: Structural and Dynamical Insights into Standard-Mechanism Protease Inhibitors. Biochemistry 58, 2524–2533 (2019).

68. Salameh, M. A., Soares, A. S., Hockla, A. & Radisky, E. S. Structural basis for accelerated cleavage of bovine pancreatic trypsin inhibitor (BPTI) by human mesotrypsin. Journal of Biological Chemistry 283, 4115–4123 (2008).

69. Peräkylä, M. & Kollman, P. A. Why does trypsin cleave BPTI so slowly? J. Am. Chem. Soc. 122, 3436–3444 (2000).

70. Yamamoto+, T., et al. Molecular Cloning and Nucleotide Sequence of Human Pancreatic Secretory Trypsin Inhibitor (PSTI) cDNA. Biochem. Biophys. Res. Commun. 132, 605–612 (1985).

71. Nagel, F. et al. Structural and Biophysical Insights into SPINK1 Bound to Human Cationic Trypsin. Int. J. Mol. Sci. 23, (2022).

72. Fraser, B. J. et al. Structure and activity of human TMPRSS2 protease implicated in SARS-CoV-2 activation. Nat. Chem. Biol. 18, 963–971 (2022).

73. Fraser, B. et al. Structural basis of TMPRSS11D specificity and autocleavage activation. Nat. Commun. (2025).

74. Pendlebury, D. et al. Sequence and conformational specificity in substrate recognition: Several human Kunitz protease inhibitor domains are specific substrates of mesotrypsin. Journal of Biological Chemistry 289, 32783–32797 (2014).

75. Yuan, C. et al. Structure of catalytic domain of Matriptase in complex with Sunflower trypsin inhibitor-1. BMC Struct. Biol. 11, (2011).

76. Boy, R. G. et al. Sunflower trypsin inhibitor 1 derivatives as molecular scaffolds for the development of novel peptidic radiopharmaceuticals. Molecular imaging and biology : MIB : the official publication of the Academy of Molecular Imaging 12, 377–385 (2010).

77. Swedberg, J. E. et al. Highly Potent and Selective Plasmin Inhibitors Based on the Sunflower Trypsin Inhibitor-1 Scaffold Attenuate Fibrinolysis in Plasma. J. Med. Chem. 62, 552–560 (2019).

78. Stolk, J. et al. Efficacy and safety of inhaled α1-antitrypsin in patients with severe α1-antitrypsin deficiency and frequent exacerbations of COPD. European Respiratory Journal 54, (2019).

79. Gaggar, A. et al. Inhaled alpha1-proteinase inhibitor therapy in patients with cystic fibrosis. Journal of Cystic Fibrosis 15, 227–233 (2016).

80. Martin, S. L. et al. Safety and efficacy of recombinant alpha1-antitrypsin therapy in cystic fibrosis. Pediatr. Pulmonol. 41, 177–183 (2006).

81. Brand, P. et al. Lung deposition of inhaled α1-proteinase inhibitor in cystic fibrosis and α1-antitrypsin deficiency. European Respiratory Journal 34, 354–360 (2008).

82. Redondo-Calvo, F. J. et al. Inhaled aprotinin reduces viral load in mild-to-moderate inpatients with SARS-CoV-2 infection. Eur. J. Clin. Invest. 52, (2022).

83. Bianchera, A. et al. Nebulizers effectiveness on pulmonary delivery of alpha-1 antitrypsin. Drug Deliv. Transl. Res. 13, 2653–2663 (2023).

84. Afar, D. E. H. et al. Catalytic cleavage of the androgen-regulated TMPRSS2 protease results in its secretion by prostate and prostate cancer epithelia. Cancer Res. 61, 1686–1692 (2001).

85. Holt-Danborg, L. et al. Insights into the regulation of the matriptase-prostasin proteolytic system. Biochemical Journal 477, 4349–4365 (2020).

86. Tamir, A. et al. The serine protease prostasin (PRSS8) is a potential biomarker for early detection of ovarian cancer. J. Ovarian Res. 9, (2016).

87. Wang, H. et al. Structure-based discovery of dual pathway inhibitors for SARS-CoV-2 entry. Nat. Commun. 14, (2023).

88. Shapira, T. et al. A TMPRSS2 inhibitor acts as a pan-SARS-CoV-2 prophylactic and therapeutic. Nature 605, 340–348 (2022).

89. Wettstein, L. et al. Peptidomimetic inhibitors of TMPRSS2 block SARS-CoV-2 infection in cell culture. *Commun*. Biol. 5, (2022).

90. Papa, G., Id, D. L. M., Albecka, A. & Id, L. G. W. Furin cleavage of SARS-CoV-2 Spike promotes but is not essential for infection and cell-cell fusion. PLoS Pathog. 17, 1–20 (2021).

91. Barash, S., Wang, W. & Shi, Y. Human secretory signal peptide description by hidden Markov model and generation of a strong artificial signal peptide for secreted protein expression. Biochem. Biophys. Res. Commun. 294, 835–842 (2002).

92. Wu, T. et al. Three Essential Resources to Improve Differential Scanning Fluorimetry (DSF) Experiments. BioRxiv https://www.biorxiv.org/content/10.1101/2020.03.22.002543v1.full (2020).

