## Supplementary Information for "Hepatocyte growth factor activator inhibitor-2 rapidly inactivates airway-expressed human Type II Transmembrane Serine Proteases"

**for**

*Corresponding authors

**Figures**

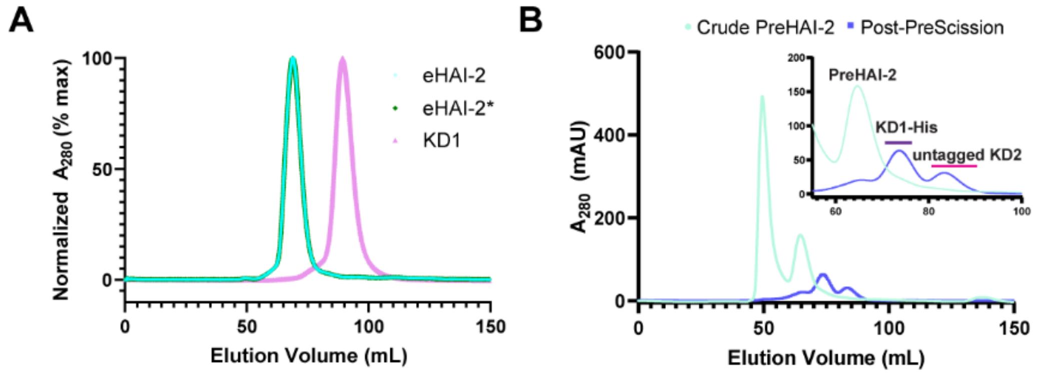


**Fig. S1. Size-exclusion chromatography separation of HAI-2 and PreHAI-2 proteins.** **A**, Size-exclusion chromatography chromatograms of the purified eHAI-2-His (teal), eHAI-2*-His (green) and KD1-His (pink) proteins. All proteins were loaded to a HiLoad 16/60 Superdex 75 gel filtration column. Protein content, monitored by $A_{280}$, was normalized to the maximum observed signal and plotted as a percentage. **B**, Size-exclusion chromatography profiles of crude PreHAI-2 (teal) overlaid with purified PreHAI-2 treated with PreScission Protease for 38 hours (blue) to facilitate untagged KD2 domain release. Bars indicate the fractions pooled for isolation of untagged KD2 (pink) and TEV-His-KD1 (purple).

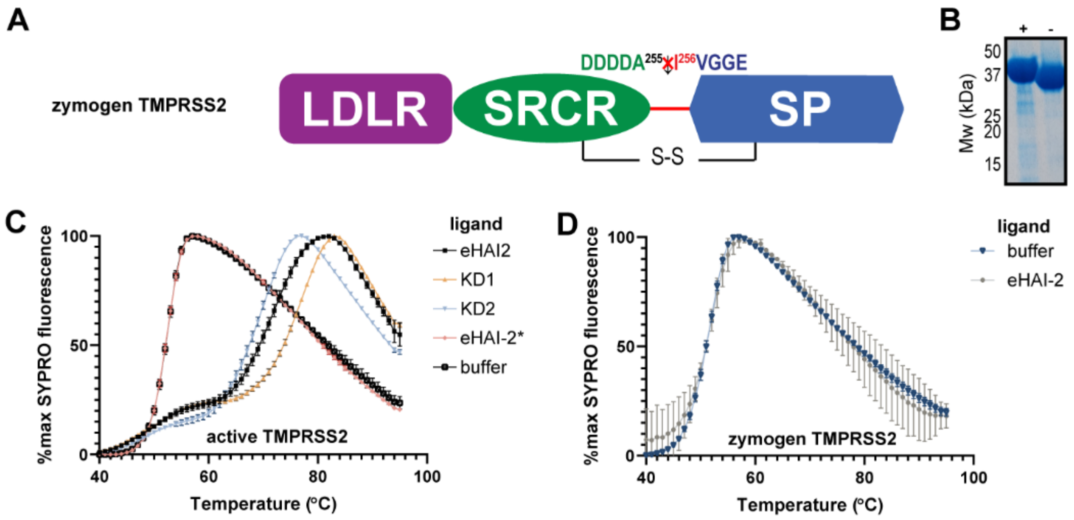


**Fig. S2. Design of a zymogen-locked TMPRSS2 protein and DSF ligand binding assays with zymogen and active TMPRSS2.** **A**, Schematic of the zymogen TMPRSS2 protein construct with a DDDDA255 amino acid substitution within the TMPRSS2 zymogen activation motif. **B**, SDS-PAGE analysis of the purified zymogen TMPRSS2 protein. SDS-PAGE samples were thermally denatured and reduced (5 mM 2-mercaptoethanol, 5 min 95 C; indicated with +) or were not thermally denatured and reduced (indicated with -) prior to gel separation. **C**, TMPRSS2 melt curves in the presence of the indicated HAI-2 protein or vector control (buffer), detected by fluorescence (excitation:emission of 465:580) with the fluorogenic SYPRO orange dye. Approximately 2 μg active TMPRSS2 and 5X SYPRO orange were used per well. **D**, Zymogen TMPRSS2 melt curves in the presence of eHAI-2 or vector control (buffer).

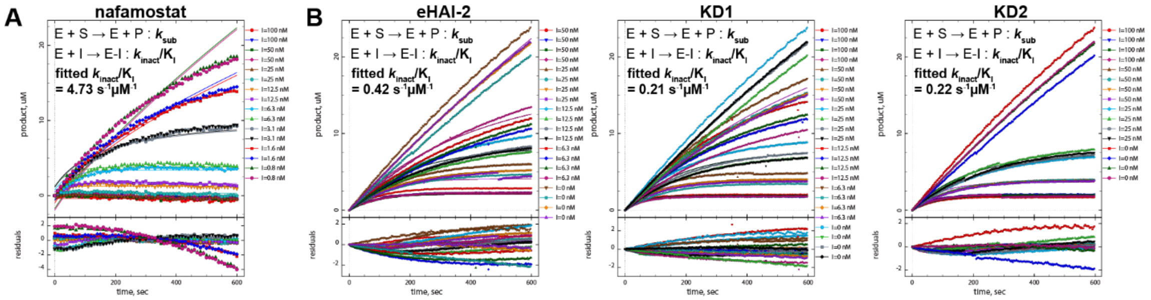


**Fig. S3. TMPRSS2 inactivation kinetics with nafamostat and HAI-2 proteins.** **A**, Kinetic model and curve-fitted data for TMPRSS2 incubated with the indicated concentrations of nafamostat or (**B**) HAI-2 proteins and 100 µM Boc-QAR-AMC substrate. Model parameters for curve fitting are shown, with residual values from the data curve fitting shown below the plot.

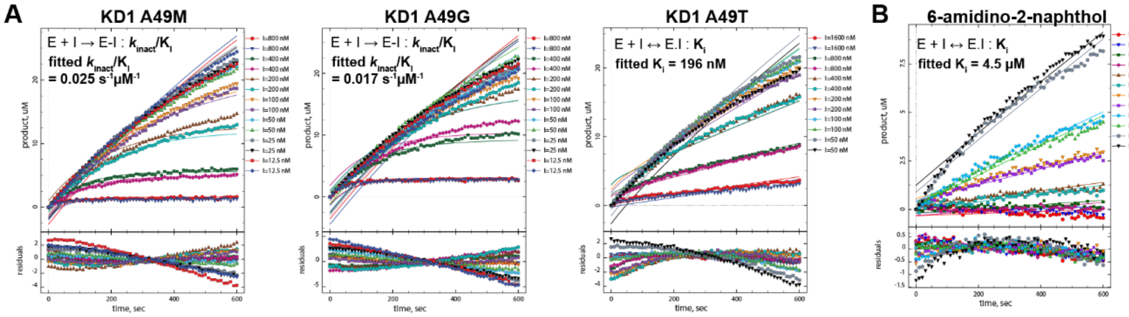


**Fig. S4. TMPRSS2 inactivation and inhibition kinetics of HAI-2 KD1 mutant proteins. A**, Kinetic model and curve-fitted data for TMPRSS2 incubated with the indicated concentrations of KD1 A49M, A49G, and A49T mutant proteins and 100 µM Boc-QAR-AMC substrate. **B**, Kinetic model and curve-fitted data for TMPRSS2 incubated with the indicated concentrations of 6-amidino-2-naphthol.

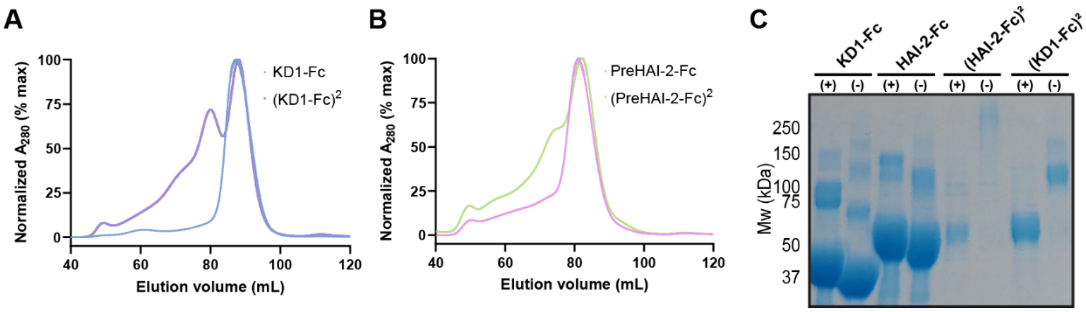


**Fig. S5. Purification of Fc-tagged HAI-2 proteins secreted from Expi293F cells. A**, Size-exclusion chromatography (SEC) purification of the KD1-Fc (teal) and (KD1-Fc)$^2$ (violet) proteins. Proteins were loaded to a HiLoad 16/60 Superdex 75 gel filtration column. Protein content, monitored by $A_{280}$, was normalized to the maximum observed signal and plotted as a percentage. **B**, SEC purification of the eHAI-2-Fc (pink) and (eHAI-Fc)$^2$ (lime) proteins. **C**, SDS-PAGE analysis of purified, Fc-tagged HAI-2 proteins. Prior to SDS-PAGE separation, samples were thermally denatured and reduced (+) or not thermally denatured and not reduced (-).

### Tables

**Table S1. Ligation independent cloning (LIC) mutagenesis primer sequences for recombinant HAI-2 and TMPRSS2 protein constructs.** All constructs contain a N-terminal honeybee mellitin signal sequence and a C-terminal EFVEHHHHHHHH affinity tag, unless otherwise indicated.

| **Protein construct (residues)** | | **Primer sequence (5' to 3')** |
|---|---|---|
| eHAI-2 (29-197) | Fwd | tacatctatgcggccgctgaccgagaacgcagcatcc |
| | Rev | ggtgctcgacgaattcctttgagccaaggggcagg |
| HAI-2 KD1 (29-91) | Fwd | tacatctatgcggccgctgaccgagaacgcagcatcc |
| | Rev | ggtgctcgacgaattcgacagtggcacatttcttgag |
| HAI-2 KD2 (130-186) | Fwd | tacatctatgcggccgctgaagaatactgcaccgccaac |
| | Rev | ggtgctcgacgaattcctggcggaagcagcggag |
| eHAI-2* (29-197) | Gene synthesized | |
| PreHAI-2† | Fwd (1;2) | ttgtatttccagggcgaccgagaacgcagcatcc;<br>agcaggctcgaagttcttttccagggaccaagtgctcccagaaggcagg |
| | Rev (1;2) | acttggtccctggaaaagaacttcgagcctgctggtggccaggtcacc;<br>caagcttcgtcatcactttgagccaaggggcagg |
| HAI-2 KD1 A49G (29-91) | Fwd | tacatctatgcggccgctgaccgagaacgcagcatcc |
| | Rev | ggtgctcgacgaattcgacagtggcacatttcttgag |
| HAI-2 KD1 A49S (29-91) | Fwd (1;2) | tacatctatgcggccgctgaccgagaacgcagcatcc;<br>tgccggagctccatgcctaggtggtggtac |
| | Rev (1;2) | catggagctccggcatctgcccaccaccttcg;<br>ggtgctcgacgaattcgacagtggcacatttcttgag |
| HAI-2 KD1 A49M (29-91) | Fwd | tacatctatgcggccgctgaccgagaacgcagcatcc |
| | Rev | ggtgctcgacgaattcgacagtggcacatttcttgag |
| HAI-2 KD1 A49T (29-91) | Fwd | tacatctatgcggccgctgaccgagaacgcagcatcc |
| | Rev | ggtgctcgacgaattcgacagtggcacatttcttgag |
| Zymogen TMPRSS2 (109-492)† | Fwd (1;2;3) | tacatctatgcggccgctatgagcggcctgaacg;<br>attgaatggcatgaaatgggcagcaagtgctccaac;<br>gacgacgcgatcgtgggcggcgagagcgcgctc |
| | Rev (1;2;3) | ttcatgccattcaattttctgcg;<br>acgatcgcgtcgtcgtcgtctgagttcaagttga;<br>ggtgctcgacgaattcgccgtctgccctcatttgtcg |
